# The histone chaperone activity of SPT2 controls chromatin structure and function in Metazoa

**DOI:** 10.1101/2023.02.16.528451

**Authors:** Giulia Saredi, Francesco N. Carelli, Giulia Furlan, Stephane Rolland, Sandra Piquet, Alex Appert, Luis Sanchez-Pulido, Jonathan L. Price, Pablo Alcon, Lisa Lampersberger, Anne-Cécile Déclais, Navin B. Ramakrishna, Rachel Toth, Chris P. Ponting, Sophie E. Polo, Eric A. Miska, Julie Ahringer, Anton Gartner, John Rouse

**Author notes:** These authors contributed equally: Francesco N. Carelli, Giulia Furlan, Stephane Rolland, Sandra Piquet, Alex Appert. Transine Therapeutics, Babraham Hall, Cambridge CB22 3AT, UK.

## Abstract

Histone chaperones control nucleosome density and chromatin structure. In yeast, the H3-H4 chaperone Spt2 controls histone deposition at active genes but its roles in metazoan chromatin structure and organismal physiology are not known. Here we identify the *Caenorhabditis elegans* orthologue of SPT2 (CeSPT-2) and show that its ability to bind histones H3-H4 is important for germline development and transgenerational epigenetic gene silencing, and that *spt-2* mutants display signatures of a global stress response. Genome-wide profiling showed that CeSPT-2 binds to a range of highly expressed genes, and we find that *spt-*2 mutants have increased chromatin accessibility at these loci. We also show that human SPT2 controls the incorporation of new H3.3 into chromatin. Our work reveals roles for SPT2 in controlling chromatin structure and function in Metazoa.

## Introduction

Genomic DNA is packaged into chromatin with the help of histone proteins. The basic units of chromatin are nucleosomes, consisting of 146 bp of DNA wrapped around a histone octamer. The octamer comprises a core histone H3-H4 tetramer flanked by two histone H2A-H2B dimers^1^. The density of nucleosomes in any given region of the genome controls the accessibility of genomic DNA to proteins involved in DNA replication, transcription and DNA repair; local nucleosome density must be altered to facilitate these key processes and restored afterwards^2-4^. A range of proteins in cell nuclei control nucleosome density and composition, including histone chaperones, nucleosome remodelling complexes, histone readers, and histone-modifying enzymes.

Histone chaperones are a structurally diverse class of proteins that bind histones and regulate nucleosome assembly and composition through a variety of mechanisms. These include shuttling of histones between cytoplasm and nucleoplasm, modulating histone stability, and facilitating histone eviction and deposition within nucleosomes^5,6^. Histone chaperones are usually specific for either histones H3-H4, or histone H2A-H2B, and sometimes display specificity for histone variants and their association can be regulated by histone post-translational modifications (PTMs)^5-7^. From a structural perspective, histone chaperone binding shields functional histone interfaces, such as histone-DNA interaction surfaces and histone dimerization domains, that are otherwise engaged when histones are assembled into nucleosomes^5^. Because of their central role in histone metabolism, histone chaperones play crucial roles in DNA replication, transcription and DNA repair^3,5,8^.

Transcription has a major impact on chromatin structure, especially as the progression of the RNA polymerase II (RNAPII) complex disrupts nucleosomes when it encounters them in the DNA template, leading to “unpeeling” of DNA from the histone octamer^9-11^. Cryo-EM studies revealed that, upon engaging with a nucleosome, RNAPII complexes stall at specific locations along the “unpeeled” nucleosomal DNA^11,12^. During this process, the histone surfaces bound by nucleosomal DNA are transiently exposed and recognised by histone chaperones^13-15^. For example, the histone chaperones SPT6, SPT5, ASF1 and the HIRA and FACT chaperone complexes have been implicated in promoting histone disassembly and recycling at active genes^8,16-19^. Conditional depletion of yeast FACT subunits or Spt6 leads to a transcription-dependent loss of H3-H4 from genes, mis-localisation of parental histone PTMs, and compensatory increase in new histone deposition over gene bodies^17,20^. Similarly, impaired binding of yeast Spt5 to histone H3-H4 leads to reduced nucleosome occupancy at active genes, loss of H3K4 trimethylation from transcription start sites (TSS) and lethality^16^. Active genes are enriched for the histone replacement variant H3.3^21-23^. Histone H3.3-H4 deposition during transcription is mediated by the HIRA complex^21,22,24^, which is composed of three subunits: HIRA, UBN1 and CABIN1. HIRA promotes both the incorporation of new H3.3-H4 as well as the recycling of parental H3.3-H4^18^, in a manner that specifically requires HIRA interaction with the UBN1 or ASF1 histone chaperones^18^, respectively. Therefore, the preservation of chromatin structure and nucleosome density during transcription requires multiple histone chaperones with overlapping but non-equivalent functions.

Budding yeast Spt2 is a poorly understood histone chaperone implicated in histone H3-H4 recycling during transcription^25,26^. Spt2 (also called Sin1) was identified genetically: mutations in the *SPT2* gene suppress transcriptional initiation defects associated with transposon insertions in the *HIS4* gene promoter^27^ or mutations in the Swi/Snf or SAGA chromatin remodelling complexes^28^. Further work revealed that Spt2 associates with the protein-coding regions of highly expressed genes, in a manner that requires Spt6^26,29^. Moreover, loss of Spt2 results in decreased association of H3 with these regions^26^. Yeast Spt2 was shown to bind to cruciform DNA *in vitro*^30^, which is thought to reflect an affinity for crossed DNA helices, reminiscent of DNA at the entry-exit of a nucleosome^31^. Yeast cells lacking Spt2 show an increase in spurious transcription from cryptic intragenic start sites^26,27^, and Spt2 mutations in the H3-H4 binding domain recapitulate these defects^25^. Furthermore, Spt2 cooperates with the yeast Hir (HIRA) complex in suppressing spurious transcription^26^. Therefore, Spt2 plays an important role in regulating H3-H4 recycling and chromatin structure in yeast. Little is known, however, about the roles and regulation of SPT2 beyond budding yeast. Chicken (*Gallus gallus*) *SPT2* is a non-essential gene, the product of which is found in both the nucleoplasm and nucleoli^32^. Chicken SPT2 interacts with RNA polymerase I (RNAPI) and was reported to support RNAPI-mediated transcription, as measured *in vitro* by nuclear run-on assay^32^. Both the DNA binding and histone binding regions of chicken SPT2 are necessary to support this function^32^. Even though almost nothing is known about SPT2 function in human cells, an X-ray crystal structure of the histone binding domain (HBD) of human SPT2 bound to a H3-H4 tetramer has been reported^25^. This analysis revealed two alpha helices (*α*C1 and *α*C2) and a connecting loop which all make contact with the tetrameric form of H3-H4. Mutating conserved residues in *α*C2, or mutating Met641 located in the inter-helical loop, reduced SPT2 binding to H3-H4 *in vitro*^25^. Replacing the HBD in yeast Spt2 with the human HBD suppresses cryptic transcription, similar to wild type yeast Spt2, but mutating Met641 in the chimeric protein blocks this suppression^25^. These data suggest that human SPT2 can regulate H3-H4 function, at least in yeast, but similar roles in human cells have not yet been described.

In this study, we dissect the *in vivo* function of SPT2 using the model organism *C. elegans*. We combine structural modelling, biochemistry and genetics approaches to characterise how SPT2 binding to histone H3-H4 regulates chromatin structure and function in Metazoa, and we show that worm SPT2 regulates chromatin density at highly expressed genes, transgenerational epigenetic silencing, and animal fertility upon heat stress. We also provide evidence that SPT2 regulates chromatin assembly in human cells.

## Results

### Identification of a *C. elegans* orthologue of the SPT2 histone chaperone

We set out to test if SPT2 histone binding activity is relevant for chromatin structure and function in Metazoa. The nematode *C. elegans* has proven a valuable system to investigate the role of chromatin modulators at the cell and organism level^33-38^, and we decided to interrogate a role for SPT2 in this organism first. However, no *C. elegans* orthologue of SPT2 had been reported when we started this project. Iterative similarity searches revealed the uncharacterized open reading frame *T05A12*.*3* as a putative orthologue. Multiple sequence alignments defined three evolutionarily conserved regions of the T05A12.3 protein product. The first region spans residues 1-129 (red box) and is conserved in metazoan orthologues but not in budding yeast (Fig. 1a). The second region spans residues 250-276 (yellow box) and is conserved from yeast to humans. The functions of these domains are unknown. The third region, spanning residues 572 to 661, is the region of highest conservation and corresponds to the histone binding domain (HBD) found in the human and yeast Spt2 orthologues^25^ (Fig. 1a, purple box; Fig. S1a). We used the crystal structure of the human SPT2 HBD^25^ as a search template to generate a structural homology model for the corresponding region of T05A12.3, which revealed three points of similarity between the two proteins. First, the tertiary structure of the putative T05A12.3 HBD adopts an arrangement that is strikingly similar to the human HBD: αC1 and αC2 helices connected by a loop (Fig. 1b). Second, both helices and the loop contact H3-H4 in our model, and the residues involved are conserved. For example, Glu637 and Glu638 in the worm T05A12.3 HDB correspond to Glu651 and Glu652 in the αC2 helix of the human HBD known to be required for H3-H4 binding^25^. Also, Met627 in the T05A12.3 HBD contacts H4 in our model; Met627 is the equivalent of Met641 in the human protein which contacts H4 and contributes to H3-H4 binding^25^ (Fig. 1c, Fig. S1b). Third, the most highly conserved residues in each of the two helices and loop lie at the interface with H3-H4 (Fig. 1c). Therefore, structural modelling strongly suggests that worm T05A12.3 is a H3-H4 binding orthologue of SPT2, and we refer to T05A12.3 hereafter as CeSPT-2.

**Figure 1.**
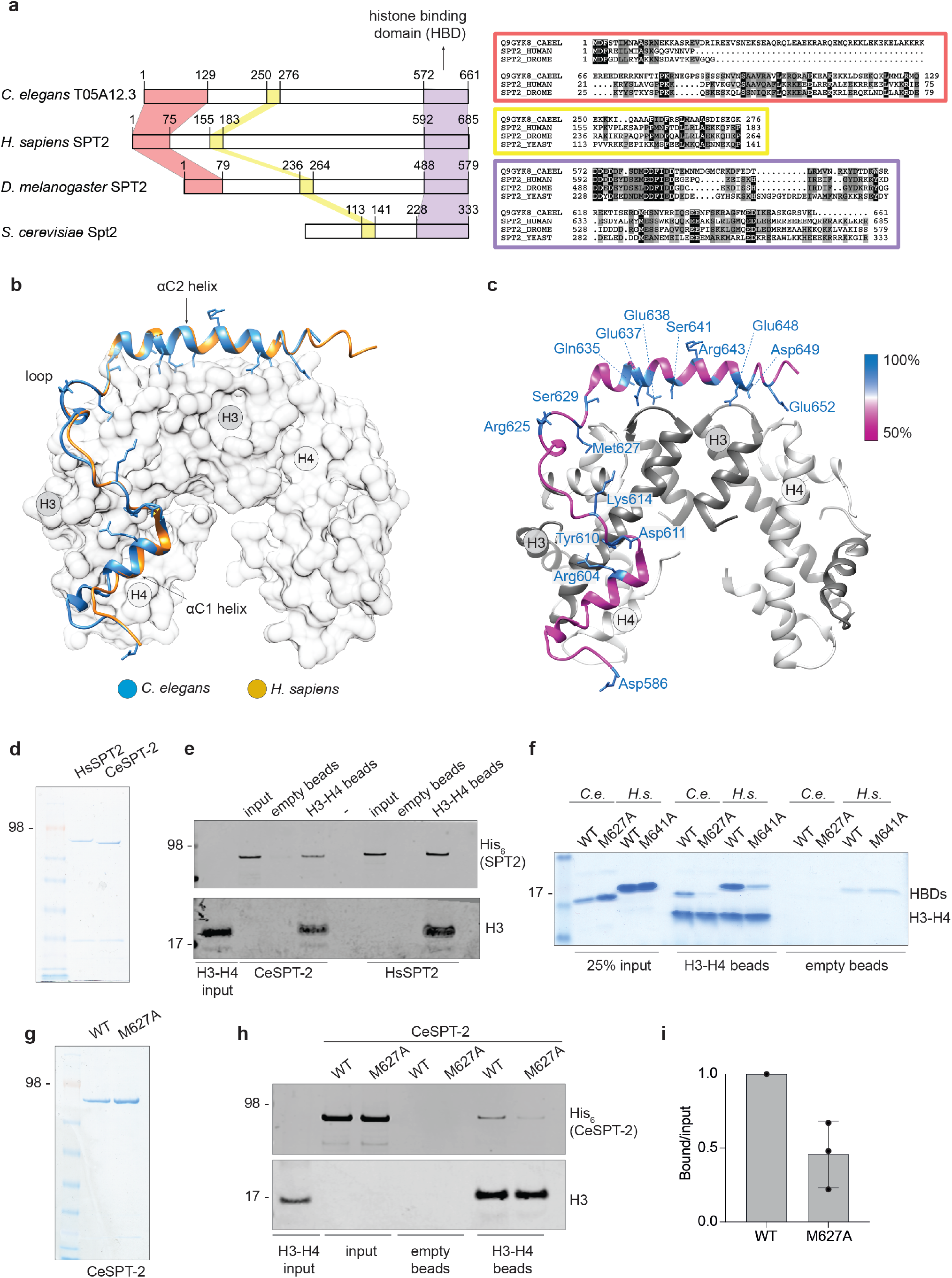
Identification of the *C. elegans* orthologue of SPT2. **a**, Schematic representation of the three evolutionarily conserved regions found in SPT2 orthologues and in *C. elegans T05A12*.*3* (left panels). N-terminal, central and C-terminal (HBD) regions are shown shaded in red, yellow, and violet, respectively. Multiple sequence alignments corresponding to the three conserved regions are shown inside coloured boxes in red, yellow, and violet, respectively (right panels). The amino acid colouring scheme indicates the average BLOSUM62 score (correlated to amino acid conservation) in each alignment column: black (greater than 3.5), grey (between 3.5 and 1.5) and light grey (between 1.5 and 0.5). Sequences are named according to their UniProt identifier. Species abbreviations: Q9GYK8_CAEEL, *Caenorhabditis elegans*; SPT2_HUMAN, *Homo sapiens*; SPT2_DROME, *Drosophila melanogaster*; SPT2_YEAST; *Saccharomyces cerevisiae*. **b**, Structural homology model for the putative HBD of *C. elegans* T05A12.3 (blue). This was generated using the crystal structure of the *H. sapiens* SPT2 HBD (yellow) in complex with the H3-H4 tetramer (shaded white) as a search template (PDB code 5BS7^25^). The positions of the *α*C1 and *α*C2 helices and the connecting loop are shown. **c**, Same as d, except that only the CeSPT-2 HBD is shown, colour-coded according to the degree of amino acid conservation. **d**, Coomassie gel staining of recombinant full-length HsSPT2 and CeSPT-2 produced in bacteria. **e**, Pull-down of full-length recombinant CeSPT-2 or human HsSPT2 with beads covalently coupled to histone H3-H4 in the presence of 500mM NaCl. One representative experiment of two is shown. **f**, H3-H4 pull-down with recombinant CeSPT-2 HBD (wild type (WT) or histone binding mutant (M627A)), or HsSPT-2 HBD (wild type (WT) or histone binding mutant (M641A)). One representative experiment of three is shown. **g**, Coomassie gel staining of recombinant full-length CeSPT-2 (WT and M627A). **h**, H3-H4 pull-down with full-length recombinant CeSPT-2 WT and M627A. **i**, Quantification of the three independent replicates of the H3-H4 pull-downs. n=3; data are represented as mean ± S.D.

We next tested if CeSPT-2 binds to histone H3-H4 *in vitro*. To this end we purified His_6_-tagged, full-length recombinant CeSPT-2 and human (Hs) SPT2 (Fig. 1d) and performed a pull-down experiment using recombinant histone H3-H4 complex covalently coupled to beads. As shown in Fig. 1e, CeSPT-2 binds histone H3-H4 *in vitro* similar to HsSPT2, and the isolated putative HBD of CeSPT2 also bound to H3-H4 (Fig. 1f, WT HBD). We next tested the effect of substituting Met627, the equivalent of Met641 which contacts H4 in the H3-H4 tetramer^25^ (Fig. 1c, Fig. S1b). Substituting Met627 for Ala (M627A) in the isolated CeSPT2 HBD reduced, but did not abolish, binding to immobilized H3-H4, similar to the M641A substitution in the HsSPT2 HBD analysed in parallel (Fig. 1f). Similar results were obtained using purified full-length CeSPT-2 (Fig. 1g-i). We also found that CeSPT-2 binds to synthetic cruciform DNA (Fig. S1c), similar to HsSPT2^32^ and that the CeSPT2 M627A substitution had no apparent effect on cruciform DNA binding (Fig. S1d). Hereafter, we refer to the <u>h</u>istone <u>b</u>inding-defective <u>m</u>utation encoding the CeSPT-2 M627A substitution as “HBM”.

We expected CeSPT-2 to be a nuclear protein given that it binds H3-H4 and DNA. Analysis of a worm strain in which GFP was inserted at the 5′ end of the *T05A12*.*3* gene showed that GFP-tagged CeSPT-2 is a widely expressed protein found, for example, in the head, germline, hypodermis, intestine and vulva cell nuclei (Fig. S1e, upper row). Moreover, knock-in of the HBM mutation did not affect GFP::CeSPT-2 expression or localisation (Fig. S1e, lower row). Taken together, the data above indicate that CeSPT-2 is a widely expressed, nuclear protein which appears to be the orthologue of the SPT2 histone H3-H4 chaperone.

### Loss of CeSPT-2 histone binding activity causes germline defects and temperature-dependent sterility

In order to study the impact of CeSPT-2 on worm development and fertility, two independently derived *spt-2* null strains, with the open reading frame being eliminated, were constructed, hereafter referred to as *spt-2*^KO-A^ and *spt-2*^KO-B^ (Fig. 2a). The *spt-2* null strains were viable, and their progeny size was comparable to wild type under standard growth conditions (20°C, Fig S2a). However, when worms were grown at 25°C for one generation, we noticed that a low proportion of worms produced far fewer progeny than wild type (Fig. 2b; Table S1). We next investigated if this apparent fertility defect became more pronounced in subsequent generations (Fig. 2c). As shown in Fig. 2d and e, the proportion of sterile *spt-2* null worms increased progressively with each generation, so that after 10 generations at 25°C very few or no fertile worms remained. We also generated a knock-in worm strain harbouring the HBM mutation that reduces CeSPT-2 binding to H3-H4 (*spt-2*^HBM^) (Fig. 2a). The *spt-2*^HBM^ worms also showed an increased incidence of sterility when grown at 25°C for several generations, although to a lesser extent than the *spt-2* null strains (Fig. 2e, Fig. S2b) probably because the CeSPT-2 HBM shows residual binding to H3-H4 (Fig. 1f, h). To be certain that the sterility observed in the *spt-2*^HBM^ strain is a direct consequence of M627A mutation, we reverted the Ala627 HBM mutation to wild type (Met627). The resulting strain (*spt-2*^HBM^ A627M) lost the sterility phenotype associated with *spt-2*^HBM^ indicating that the sterility is due to loss of CeSPT-2 histone binding capacity (Fig. S2b).

**Figure 2.**
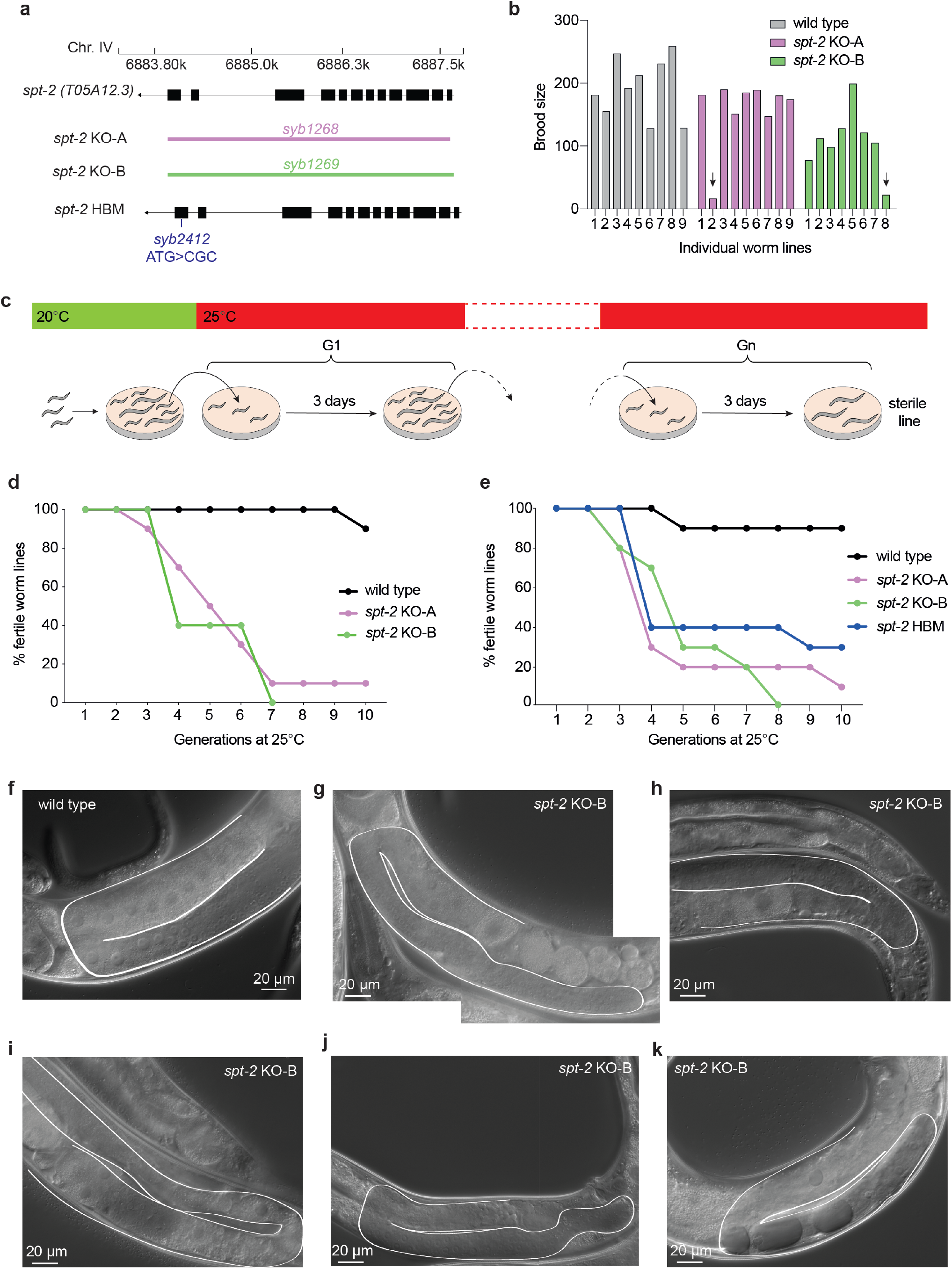
Loss of CeSPT-2 histone binding causes germline defects and sterility. **a**, Schematic diagram of the wild type, *spt-2*^KO^ and *spt-2*^HBM^ alleles. The genomic location of the *spt-2* (*T05A12*.*3*) gene on chromosome IV is shown. Coloured bars indicate the deleted regions in the *spt-2* gene in the knockout strains. **b**, The brood size of worms from the indicated strains, grown at 25°C from the L4 stage, is shown. Arrows indicate worms with low brood size. The number of worms used is indicated in the figure. **c**-**e**, Transgenerational sterility assay. Three L4 stage worms of the indicated genotypes were shifted to 25°C and grown at that temperature for the indicated number of generations. Every generation (3 days), three L3-L4 worms were moved to a new plate. A worm line was considered sterile when no progeny was found on the plate. Ten plates per genotype were used, n=10. **f-k**. Single wild type and *spt-2*^*KO-B*^ L4 worms were shifted to 25°C and grown at that temperature for the indicated number of generations; every generation (3 days), one L4 worm was moved to a new plate and the number of progeny was assessed, as indicated in Fig. S2d. Siblings of the worms that, three days after the L4 stage, showed reduced or no progeny were subjected to microscopic observation one day after the L4 stage.

To investigate the morphology of the germline in *spt-2*-defective worms, we grew *spt-*2^KO-B^ worms for several generations at 25°C and subjected the siblings of worms that showed a reduced number of progeny or sterility to microscopic analysis (Fig. S2c, d). In wild type nematode gonads, germ cells transit in an orderly manner from the mitotic stem cell state to the various stages of meiotic prophase (pachythene and diplotene) and eventually become fully mature oocytes (diakinesis stage)^39^ (Fig. 2f). In contrast, the germlines of *spt-2* null worms whose siblings had a decreased progeny size show a wide array of defects at 25°C (generations G4 and G5, Fig. S2d), including: i) a low number of fertilized embryos, which appeared rounded and incapable of hatching or further development (Fig. 2g); ii) oocytes but no productive fertilization, and presenting an abrupt transition between pachytene cells and oocytes (Fig. 2h); iii) a highly distorted germline showing mis-localization of oocytes (Fig. 2i); and iv) signs of masculinization, as evidenced by an excessive number of sperm cells (Fig. 2j). The germlines of siblings of worms that were completely sterile were reduced in volume, contained a small number of enlarged mitotic germ cells, and showed vacuolization (Fig. 2k, S2e). Taken together these data show that the onset of sterility in *spt-2* defective worms is associated with pleiotropic defects in germ line development.

### CeSPT-2 histone binding activity is required for transgenerational maintenance of nuclear RNAi-mediated gene silencing

The sterility seen in *spt-2* defective worms after several generations at elevated temperature was reminiscent of worms harbouring defects in nuclear RNA interference (RNAi)^40,41^. Nuclear RNAi is a pathway in which small double-stranded (ds) RNAs trigger heritable gene silencing^42,43^. The establishment of silencing involves dsRNA-mediated dicing of the target mRNA, while the maintenance and transgenerational inheritance of the silenced state involves the RNA-dependent synthesis of secondary siRNAs and changes in chromatin state in the RNAi target gene(s), such as histone H3 methylation at Lys9 and Lys23^44-47^. In prevailing models, the Argonaute family protein HRDE-1 binds secondary small RNAs and is required for the transgenerational inheritance of gene silencing after the dsRNA trigger has been removed. HRDE-1 directs H3K9 trimethylation at RNAi target loci, although how chromatin promotes the inheritance of gene silencing across several generations is poorly understood^48^.

We investigated a role for CeSPT-2 in the nuclear RNAi pathway, using a reporter strain we described previously^42^. In this system, a *gfp::h2b* single-copy transgene which is constitutively expressed in the worm germline can be silenced by feeding worms with bacteria expressing double-stranded *gfp* RNA (*gfp* RNAi)^42^ (Fig. 3a). Analysis of GFP fluorescence showed that silencing of the *gfp::h2b* reporter transgene occurs normally in *spt-2* null and *hrde-1*-defective worms grown on bacteria expressing the *gfp* RNAi (P0) (Figs. 3b, c). After removing the bacteria, silencing was maintained for 5 generations in the wild type worms, but the transgene was de-silenced in the first generation (G1) of *hrde-1*-defective worms. Strikingly, the reporter transgene was robustly de-silenced in both of the *spt-2* null strains from the second generation (G2) after the removal of the *gfp* RNAi, as measured by GFP fluorescence (Fig. 3b, c) and mRNA abundance (qPCR) (Fig. 3d). We also found that the *spt-2*^*HBM*^ strain showed transgene de-silencing albeit slightly later than in the null strains (Figs. 3e, f). Taken together, these data show that CeSPT-2 histone binding activity is required for the transgenerational inheritance of epigenetic gene silencing.

**Figure 3.**
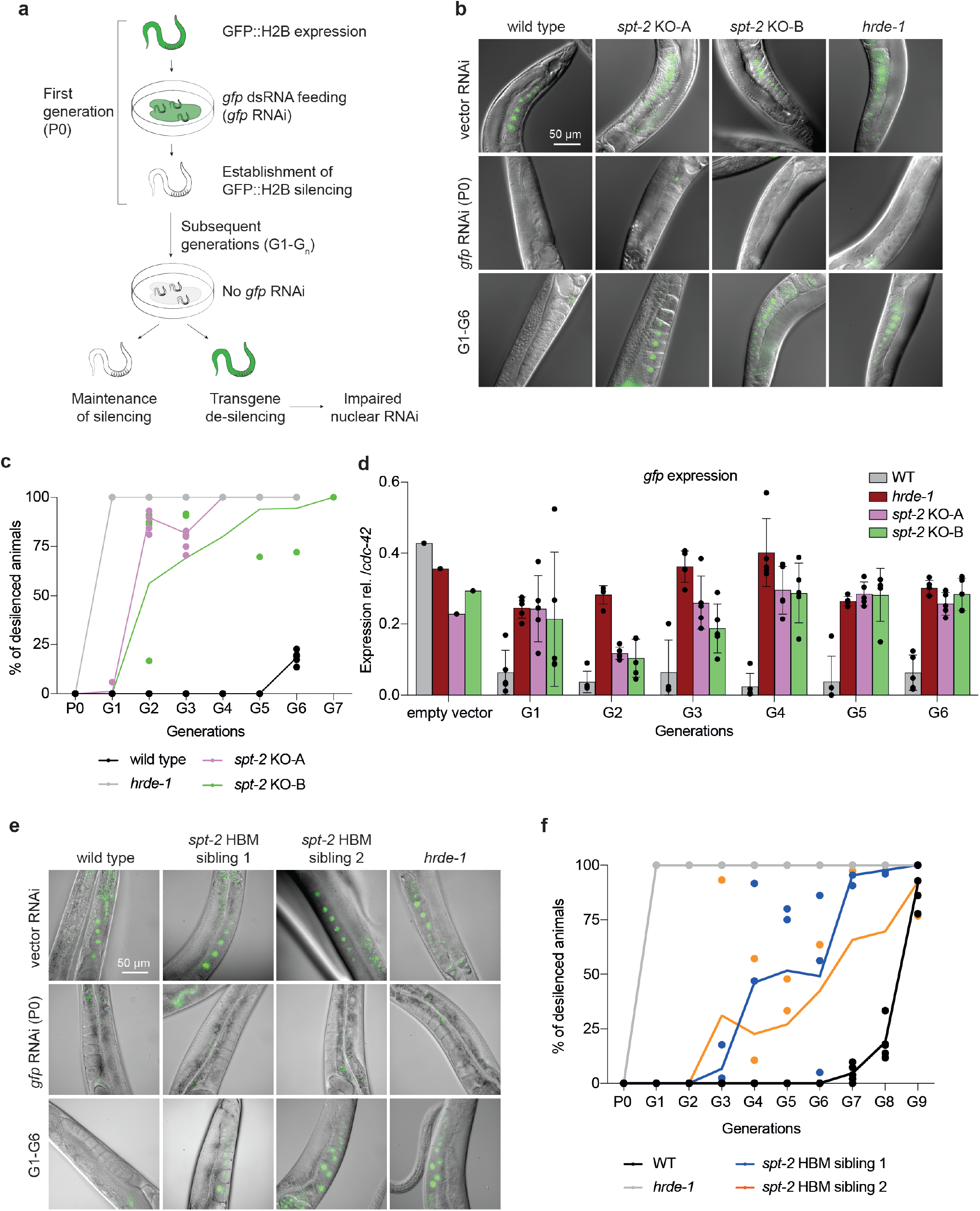
CeSPT-2 histone binding is required for transgenerational gene silencing. **a**, Schematic diagram of the transgenerational RNAi assay described previously^42^. **b**, Representative images of the germline of the *gfp::h2b* reporter worm strains without RNAi (vector RNAi), after *gfp::h2b* RNAi treatment and in the subsequent generations after removal of the dsRNA (G1-G6). **c**, Quantification of GFP::H2B expression. Worms showing “dim” GFP intensity during microscopy inspection were considered as silenced. The graph indicates the mean percentage of de-silenced worm lines across the replicate lines (mean ± S.D.), and the percentage of de-silenced worms in each replicate is indicated by a dot; n=5. **d**, *gfp::h2b* mRNA expression measured by qPCR, normalized relative to *cdc-42*. n=1 for ‘empty vector’ point; n=5 for G1-G6; data are represented as mean ± S.D. **e**, Representative microscopy images of the germline of the indicated worm strains without RNAi (vector RNAi), after *gfp::h2b* RNAi treatment and in the subsequent generations after removal of the dsRNA (G1-G9). **f**, Quantification of GFP::H2B expression, as in c. The graph indicates the mean percentage of de-silenced worm lines across the replicate lines (mean ± S.D.), and the percentage of de-silenced worms in each replicate is indicated by a dot; n=5 (WT and *hrde-1* mutant); n=3 (for each of the *spt-2*^HBM^ siblings).

### CeSPT-2 binds and controls the chromatin structure of highly transcribed genes

To gain insight into the endogenous genes that might be regulated by CeSPT-2, we sought to identify its chromatin occupancy genome-wide. We performed chromatin immunoprecipitation and sequencing (ChIP-seq) in synchronized adult worms expressing endogenously tagged GFP::CeSPT-2, using wild type worms as control. Around 88% of the genomic CeSPT-2 target sites we identified lie within genic regions (5299/6003 sites), with the remaining sites found in intergenic (∼7%, 258 sites) or repetitive (∼4%, 446 sites) sequences (Fig. 4a). GFP::CeSPT-2 associates over the entire length of what we hereby call as ‘CeSPT-2 target genes’, with an apparent enrichment for the 3’ end of the gene (Fig. 4b). We reasoned that a histone chaperone with a potential function during transcription would associate with genes in a transcription-dependent manner. Indeed, CeSPT-2 bound genes are highly expressed, and their transcriptional levels are positively correlated with CeSPT-2 chromatin binding (Fig. 4c).

**Figure 4.**
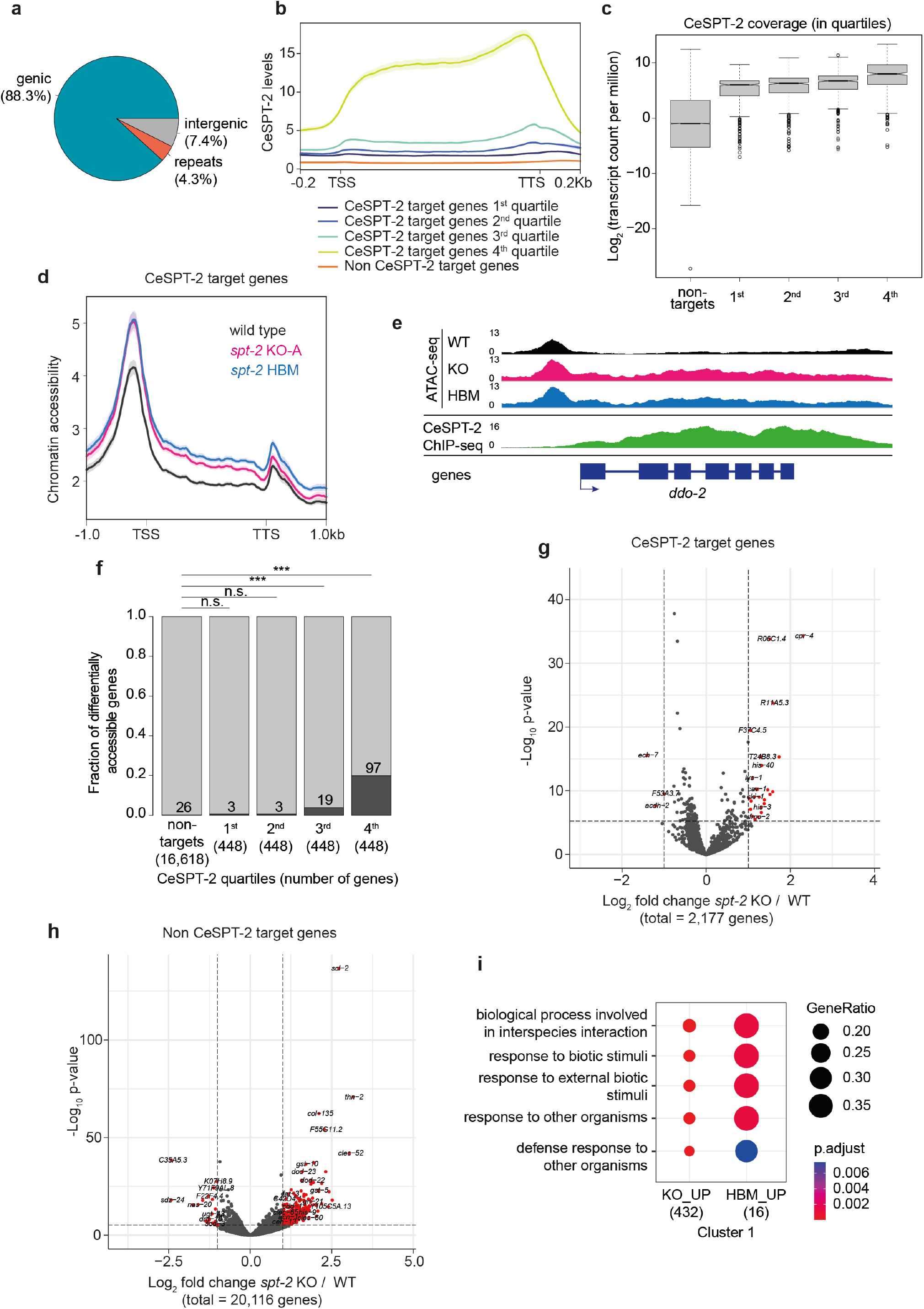
CeSPT-2 histone binding controls chromatin accessibility. **a**, Distribution of GFP::CeSPT-2 ChIP-seq peaks in different genome locations. **b**, Metagene profile of GFP::CeSPT-2 occupancy levels within the coding region of its target genes, compared to non CeSPT-2 target genes. **c**, GFP::CeSPT-2 chromatin occupancy regions were divided in quartiles, and gene expression levels (measured in synchronised wild type adult worms) were plotted in each quartile, and compared to CeSPT-2 non-target genes. The box represents the interquartile range, the whiskers the min and max (excluding outliers). **d**, Chromatin accessibility at CeSPT-2 target genes in *spt-2*^KO-A^ and *spt-2*^HBM^ adult worms, measured by ATAC-seq and expressed in reads per million. TSS, transcription start site; TTS, transcription termination site. Two independent replicates of the ATAC-seq experiment were performed. **e**, Example of ATAC-seq and GFP::CeSPT-2 ChIP-seq tracks. The ATAC-seq signal across the gene body of *ddo-2* in wild type, *spt-2*^KO-A^ and *spt-2*^HBM^ worms is indicated, with the GFP::CeSPT-2 binding profile shown in the bottom profile. **f**, Fraction of genes with significantly increased accessibility in *spt-2*^KO-A^ (dark grey) among CeSPT-2 non-target and target genes. CeSPT-2 targets were divided in quartiles based on their CeSPT-2 enrichment. Statistical difference between CeSPT-2 targets and non-targets was measured by a chi-squared test. Benjamini-Hochberg corrected p values: n.s.: p > 0.05; ***: p < 0.001. **g, h**, Volcano plot of gene expression levels of CeSPT-2 target genes (g) and non-targets (h) in *spt-2* null versus wild type worms. Red points indicate genes with FDR<0.001 and log_2_ fold change >1 or <-1. **i**, Gene Ontology (GO) analysis of the genes upregulated in the indicated *spt-2* mutants.

We next asked whether loss of CeSPT-2 – or loss of its histone H3-H4 binding activity – affects chromatin accessibility of the genes it is enriched at. We performed ATAC-seq in wild type, *spt-2*^KO-A^ and *spt-2*^HBM^ adult worms and found that chromatin accessibility increased across the entire length of the gene body of CeSPT-2 bound genes (Fig. 4d, e, Fig. S3a). In contrast, no significant increase in chromatin accessibility was observed at genomic regions that are not enriched for CeSPT-2 (Fig. S3b). Notably, most genes showing a significant increase in their accessibility in *spt-2* mutants had high levels of CeSPT-2 binding (Fig. 4f). Therefore, CeSPT-2 binds and influence the chromatin architecture of some of the most highly transcribed genes in *C. elegans*.

We next sought to evaluate a potential role for CeSPT-2 on gene expression. The expression of most CeSPT-2 bound genes did not change significantly in *spt-2* null worms compared with wild type (Fig. 4g), with only 1.8% (40/2177) and 0.18% (4/2177) targets showing a marked up- or downregulation in the mutant, respectively (FDR<0.001 and log2 fold change >1 or <-1; Supplementary Table 2). While we might have expected more accessible chromatin to result in increased gene expression, CeSPT-2 bound genes are among the most highly expressed genes in adult worms (Fig. 4c) and we speculate that a further increases in the expression of these genes may not be possible. While CeSPT-2 target gene expression appeared to be unaffected in *spt-2* null worms, global mRNA-seq analysis of CeSPT-2 non-target genes revealed a pronounced up-regulation of gene expression in these worms compared with wild type (Fig. 4h). Specifically, we found 40 protein-coding genes downregulated and 605 genes upregulated in *spt-2* null worms (FDR<0.001 and log_2_ fold change >1 or <-1; Fig. S3c: validation by qPCR using independently harvested RNA). Of the 605 up-regulated genes, only 40 were CeSPT-2 targets. The up-regulated genes were strongly enriched Gene Ontology (GO) terms related to ‘defense response’, ‘protein dimerization’ and ‘nucleosome’ (Fig. 4i, S3d), indicating that *spt-2* KO worms experience activation of a global stress response. The HBM mutants showed a similar increased expression of ‘defense response’ genes (Fig. 4i), although gene up-regulation was less dramatic in *spt-2*^HBM^ worms; around 80% of the genes up-regulated in *spt-2*^HBM^ worms are also up-regulated in the *spt-2*^KO^ strain. Taken together, these data show that CeSPT-2 binds to a range of highly expressed target genes, and its histone binding activity controls chromatin accessibility at these loci; loss of *spt-2* results in a global stress response.

### SPT2 controls new histone H3.3 levels and deposition in human cells

The data above show clear roles for CeSPT-2 activity in controlling chromatin maintenance in worms, and we explored a potential role in human cells. We first tested if HsSPT2 associates with chromatin. Affinity-purified antibodies raised against HsSPT2 recognised a band of approximately 75 kDa in extracts of U-2-OS cells, that was reduced in intensity by three different HsSPT2-specific siRNAs (Fig. S4a). Fractionation experiments showed that endogenous HsSPT2 is strongly enriched in the chromatin fraction of U-2-OS cells (Fig. 5a). Furthermore, fluorescence analysis of U-2-OS cells after pre-extraction of soluble proteins showed clearly that HsSPT2 tagged with GFP at either the N-terminus or C-terminus associates with chromatin (Fig. 5b). An HsSPT2 deletion fragment (aa 1-570) lacking the HBD bound chromatin similar to full-length HsSPT2, indicating that interaction with H3-H4 does not mediate chromatin association (Fig. 5b, S4b, c). Consistent with this idea, a deletion fragment corresponding to the HBD alone (aa 571-end) localized to the nucleus but was not retained on chromatin (Fig. 5b, S4b, c).

**Figure 5.**
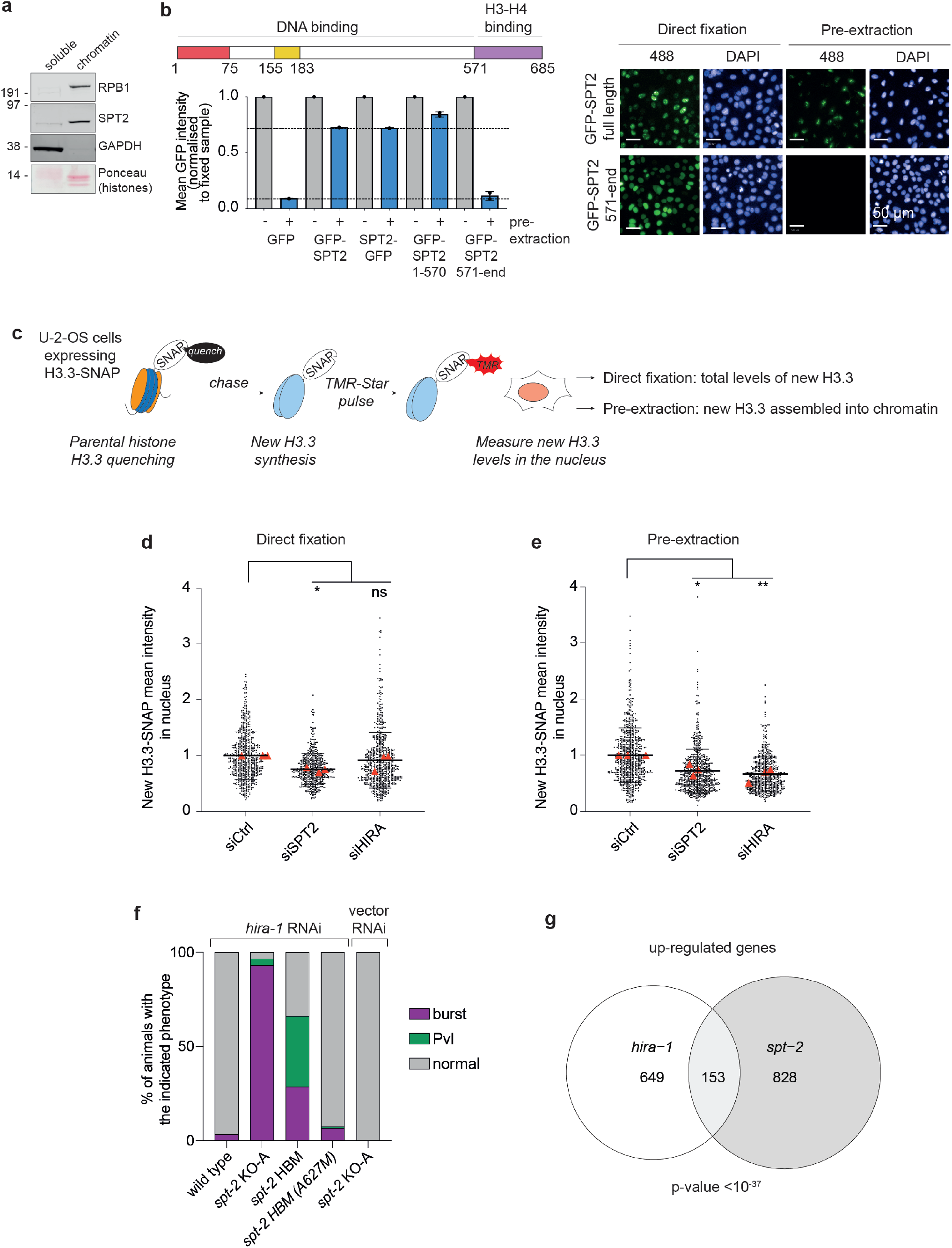
SPT2 is required for histone H3.3-H4 deposition in human cells. **a**, Distribution of HsSPT2 between soluble and chromatin fractions of U-2-OS cells. **b**, GFP-HsSPT2 binding to chromatin in fixed or pre-extracted U-2-OS cells assessed by high-content microscopy. Graph shows the mean GFP intensity normalised to the fixed sample; n=2, data are represented as mean with range of two independent experiments. **c**, Schematics of the SNAP-tag assay for new histone H3.3 incorporation. **d, e**, Fluorescence intensity of TMR-labelled SNAP-H3.3 in U-2-OS SNAP-H3.3 cell nuclei after (d) direct fixation or (e) pre-extraction. Cells were harvested 48 hours post siRNA transfection. Data are represented as mean ± S.D. from 3 independent experiments (red triangles) and normalized to the mean of siCtrl in the same experiment. **f**, Percentage of burst worms or worms showing a protruding vulva (Pvl) phenotype upon treatment with *hira-1* RNAi, or empty vector RNAi, as indicated. Forty worms were scored per replicate, and three independent replicates of the experiment were performed. **g**, Venn diagram showing the overlap between up-regulated genes in *spt-2* null and *hira-1* null worms. Hypergeometric test; enrichment: 3.085, p value < 10^−37^.

**Figure 6.**
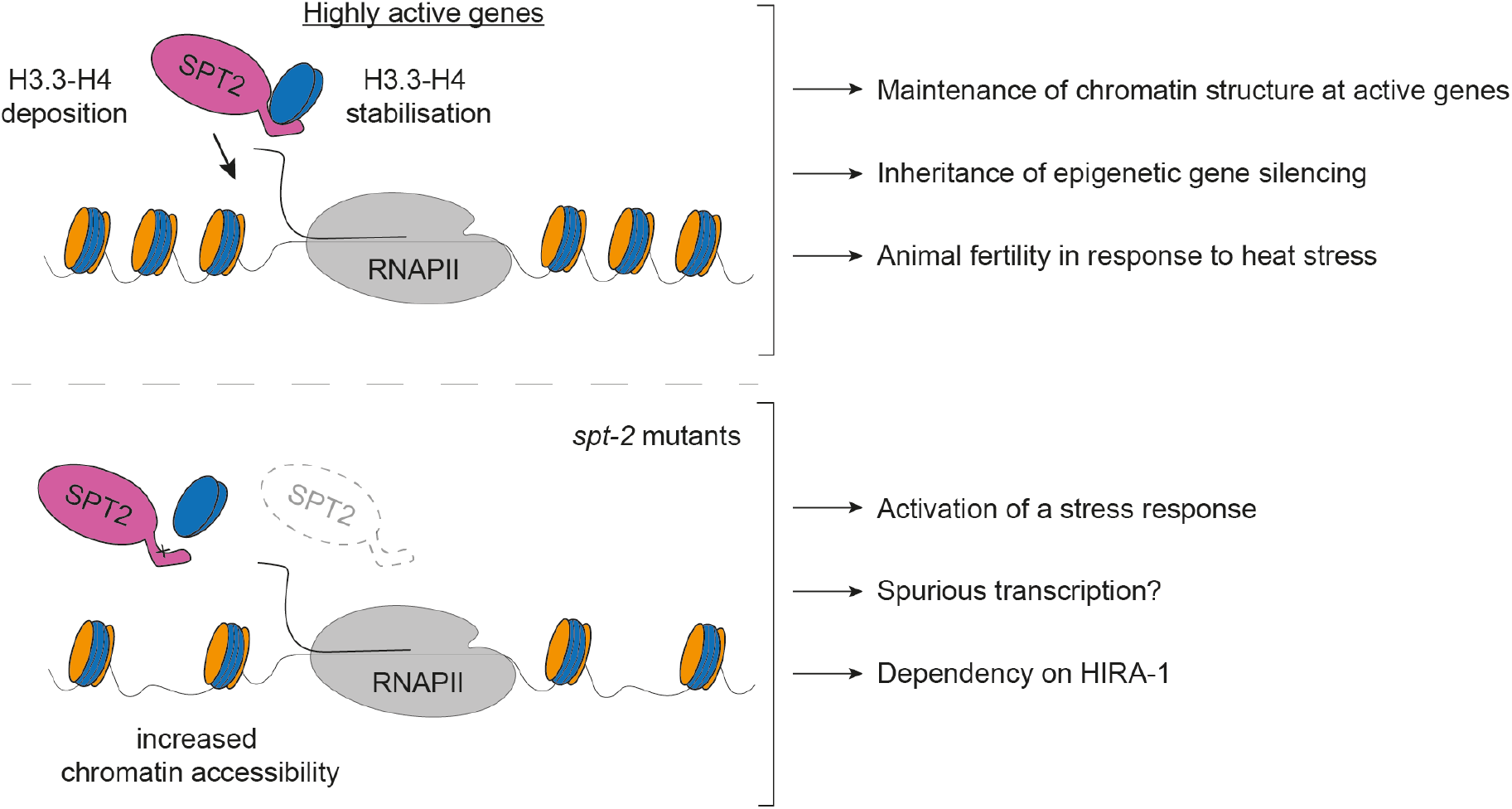
Model. SPT2 localises to highly active genes where it promotes the maintenance of chromatin structure via its histone H3-H4 binding capacity. Here, it may act via stabilising new histone H3.3 prior to their incorporation, as well as via directly promoting their incorporation into chromatin. In worms, the interaction of CeSPT2 with H3-H4 is required for the transgenerational inheritance of gene silencing and to promote the integrity of the germline under heat stress conditions. Loss of *spt-2*, or impaired SPT-2 binding to histones, activates a stress response, potentially due to aberrant transcription, and makes worms reliant on the HIRA-1 H3.3-H4 histone chaperone.

We next tested if HsSPT2 influences histone H3.3 deposition, which is enriched at active genes^22^. With this goal in mind, we employed a reporter cell line stably expressing a SNAP-tagged H3.3^49^ which can be covalently labelled *in vivo* using cell-permeable compounds to distinguish between pre-existing and newly synthetised protein pools. To specifically visualise newly incorporated H3.3, we used a quench-chase-pulse approach: we first labelled all pre-existing H3.3-SNAP with the non-fluorescent chemical bromothenylpteridine (BTP, quench), we allowed a 2-hour chase period to synthesize new histones, and we finally labelled new H3.3-SNAP with the fluorophore tetramethylrhodamine (TMR, pulse; Fig. 5c). To analyse the total levels of new H3.3-SNAP (which are a measure of H3.3 synthesis and stability), we fixed cells and measured H3.3-SNAP-TMR intensity in cell nuclei (Fig. 5c). Furthermore, to assess the incorporation of newly synthesised H3.3 into chromatin, we pre-extracted soluble proteins prior to fixing the cells and then measured TMR fluorescence intensity (Fig. 5c). As shown in Fig. 5d and e, siRNA-mediated depletion of HsSPT2 reduces both the total levels of SNAP-tagged new H3.3 (direct fixation, Fig. 5d) as well as the amount of new H3.3 incorporated into chromatin (pre-extraction, Fig. 5e). New H3.3 incorporation is reduced after depletion of HIRA (Fig. 5e), as previously shown^18,22^. We note that H3.3-SNAP mRNA levels in HsSPT2-depleted cells are comparable to control cells (Fig. S4d). Moreover, HsSPT2 knock-down does not affect HIRA stability, nor does it impair HIRA binding to chromatin (Fig. S4e, f). From these data, we conclude that human SPT2 controls histone H3.3 levels and promotes their incorporation into chromatin.

### Functional interplay between the SPT-2 and HIRA-1 histone chaperones in *C. elegans*

Given the roles for SPT2 and HIRA in new H3.3 deposition in human cells, we queried whether CeSPT-2 and HIRA-1 may overlap functionally in worms. *hira-1* null worm strains are known to be viable but show pleiotropic defects, including low brood size and morphological defects (such as small pale bodies and protruding vulvas “Pvl”)^36,37^. We set out to test the impact of the concomitant loss of CeSPT-2 and HIRA-1 on worm development and viability. With this in mind, we depleted HIRA-1 by RNAi from wild type, *spt-2* null or *spt-2*^HBM^ worms grown at 25°C. We noted that the Pvl phenotype, reported in *hira-1* null worms^37^, was not observed in wild type worms treated with *hira-1* RNAi (Fig. 5f), possibly due to partial HIRA-1 depletion. However, HIRA-1 depletion in *spt-2* null or *spt-2*^HBM^ worms resulted in a dramatic increase in the incidence of vulvar protrusions and of burst nematodes (Fig. 5f). The morphological defects of the *spt-2*^*HBM*^ were fully rescued in the reverted (A627M) strain (Fig. 5f). Comparison of the transcriptomes of *spt-2* null (this study) and *hira-1* null worms (dataset from ref.^36^) revealed a statistically significant overlap of the genes up-regulated in both mutants, the up-regulated genes being enriched for stress response transcripts (Fig. 5g). From these data, we conclude that the CeSPT-2 and HIRA-1 histone chaperones have overlapping functions in the control of H3-H4 deposition in worms, and that HIRA-1 becomes essential for worm fitness in *spt-2* mutants.

## Discussion

In this study we presented the first demonstration that SPT2 controls chromatin structure and function in Metazoa. Through bioinformatic analyses and structural modelling we identified the previously unannotated protein T05A12.3 as the worm orthologue of CeSPT-2 (Fig. 1a-c), and we found that recombinant CeSPT-2 binds H3-H4 *in vitro* in a manner similar to the yeast and human orthologues (Fig. 1d-i). Global profiling revealed that CeSPT-2 targets a range of genomic sites in worms, the vast majority (∼88%) lying within genic regions (Fig. 4a, b). Consistent with what has been observed in yeast^26^, genes enriched for CeSPT-2 are amongst the mostly highly active genes in worms (Fig. 4c), possibly reflecting a propensity of CeSPT-2 to bind accessible chromatin regions. CeSPT-2 target genes showed increased chromatin accessibility over the entire bodies of these genes in *spt-2* null and *spt-2*^HBM^ worms (Fig. 4d, e), with a direct correlation between CeSPT-2 binding levels and extent of increased accessibility (Fig. 4f); therefore CeSPT-2 activity helps preserve the integrity of chromatin in actively transcribed regions.

Intriguingly, the expression of most CeSPT-2 target genes was unaffected in the *spt-2* null and *spt-2*^HBM^ worms (Fig. 4g). However, as CeSPT-2-bound genes are among the most highly expressed in worms, it may be that their expression is already maximal in wild type worms. What, then, is the role of CeSPT-2 in limiting chromatin accessibility at these genes? Perhaps CeSPT-2 activity prevents harmful consequences of an excessively open chromatin such as spurious transcription, as observed for yeast Spt2. This might lead to the production of aberrant transcripts resembling foreign nucleic acids^50^ or neo-antigens. A wide range of stress response genes were upregulated in *spt-2* mutant worms (Fig. 4h, i), and it is tempting to speculate this protective response is triggered by aberrant transcripts. This possibility will be interesting to investigate.

The idea that CeSPT-2 binding to H3-H4 preserves chromatin integrity at highly expressed genes is supported by our demonstration that RNAi-mediated transgenerational gene silencing of a *gfp::h2b* reporter transgene requires CeSPT-2 histone binding activity (Fig. 3a-f). This finding, to our knowledge, identifies CeSPT-2 as the first histone chaperone required for nuclear RNAi in *C. elegans*. How chromatin structure controls the transgenerational inheritance of *gfp::h2b* silencing is unclear: while histone PTMs associated with silenced chromatin are observed at the RNAi target locus in response to dsRNA^44-47,51^, the SET-25 and SET-32 H3K9/K23 tri-methyltransferases are only required for maintenance of silencing in the first generation after removal of the dsRNA trigger, and dispensable afterwards^44,52^. We speculate that CeSPT-2 is recruited to the open chromatin of the active transgene, and controls silencing of the *gfp::h2b* reporter by limiting chromatin accessibility, as shown for the endogenous CeSPT-2 target genes above (Fig. 4d, f). It will be interesting in the future to test if loss of CeSPT-2 influences the repressive H3 PTMs associated with RNAi-mediated gene silencing, as well as whether other histone chaperones are required for transgenerational gene silencing.

We do not yet know how SPT2 is targeted to chromatin. The C-terminal HBD is the most highly conserved region of SPT2, but at least in human cells it is not required for association with chromatin (Fig. 5b). However, one of the other two conserved regions we identified in our bioinformatic analyses (Fig. 1a) may promote SPT2 binding to chromatin, either directly or through an as yet unidentified binding partner. We note that our sequence conservation analysis was unable to confirm the presence of a previously described HMG-box domain (InterPro: IPR009071) in SPT2^53^, which was suggested to mediate DNA binding. Examination of the SPT2 AlphaFold models also failed to reveal an L-shape arrangement of three α-helices, which is characteristic of the HMG-box domain^54^. In yeast, the first 200 amino acids of Spt2 are necessary and sufficient for Spt2 recruitment to active genes, and for association with Spt6 which recruits Spt2 to target genes^29^. Whether metazoan SPT2 binds to SPT6 (and other components of the active RNA polymerase II complex) is not known, and mechanism behind SPT2 recruitment to chromatin in Metazoa remain to be elucidated.

Besides SPT2, the histone chaperones FACT, SPT6, SPT5, ASF1, and HIRA have all been implicated in maintaining chromatin structure at active genes^17,18,20^. One possible scenario to explain the multiplicity of chaperones acting at transcribed genes is that SPT2 could work together with the other histone chaperones, potentially in a partially redundant manner, by binding different configurations of histone H3-H4 during nucleosome assembly and disassembly. Histone turnover is highest at the most highly expressed genes^55-58^, and these genes may be particularly reliant on the joint functions of CeSPT-2 together with other histone chaperones. In this light, we found that RNAi depletion of HIRA-1 causes profound defects in *spt-2* null or *spt-2*^HBM^ mutant worms, but not in wild type worms (Fig. 5f), indicating that H3-H4 binding by CeSPT-2 likely supports a step of chromatin assembly that becomes essential in the absence of HIRA-1.

Our data from human cells shows that HsSPT2 depletion decreases the total levels of new H3.3 as well as the levels of chromatin-bound new H3.3 (Fig. 5c-e), suggesting that SPT2 can also function in regulating the production and/or stability of soluble histones. We cannot exclude that the decreased incorporation of H3.3-SNAP is simply a by-product of decreased H3.3 stability although we note, for example, that mutations in histone H3 that impair binding to the MCM2 histone chaperone affect total levels of SNAP-tagged H3.1 without decreasing new H3.1 incorporation^59^. SPT2 was recently identified as an interactor of the histone chaperone and heat shock folding chaperone DNAJC9^60^; investigating a potential interplay between SPT2 and folding chaperones in promoting histone stabilisation during transcription will be of great interest.

Faithful maintenance of chromatin structure is essential to safeguard epigenetic information and cell identity, and to preserve genome stability and organism viability. Our research presents the first detailed *in vivo* characterisation of metazoan SPT2 histone chaperone function. We show the importance of CeSPT-2 in maintaining germline fertility under heat stress conditions, and its role in preserving chromatin structure at highly active genes and ensuring the transgenerational inheritance of gene silencing. Together, our work highlights the importance of understanding how the concerted action of histone chaperones come together to preserve chromatin structure and organism fitness.

## Acknowledgments

We are grateful to members of the Labib lab for invaluable advice on recombinant protein purification and *C. elegans* techniques, and to members of the Rouse lab for fruitful discussion. We thank Constance Alabert and Karim Labib for helpful comments on the manuscript; Ramasubramanian Sundaramoorthy and Tom Owen-Hughes for the gift of recombinant *Xenopus* histones; Bettina Meier and Federico Pelisch for their help with *C. elegans* genetics; Karim Labib for the gift of the His_6_-Ulp1 plasmid; Thomas Carroll for help with microscopy; and Axel Knebel for advice on protein purification. We thank the technical support of the MRC PPU including the DNA Sequencing Service, Tissue Culture team, Reagents and Services team, and the imaging platform of the Epigenetics & Cell Fate Centre (Paris); we also thank Fiona Brown and James Hastie for SPT2 antibody production and purification. GS is supported by an EMBO Long-Term Fellowship (ALTF 951-2018), a Marie Sklowdoska Curie Individual Fellowship (MSCA-IF ‘ICL-CHROM’) and a SULSA ECR Development Fund; this project has received funding from the European Union’s Horizon 2020 research and innovation programme under the Marie Skłodowska-Curie grant agreement No 845448. This work was supported by the Medical Research Council (grant number MC_UU_12016/1) and the pharmaceutical companies supporting the Division of Signal Transduction Therapy Unit (Boehinger-Ingelheim, GlaxoSmithKline, and Merck KGaA) (JR lab) and by a BBSRC grant (BB/S002782/1) to AG and JR. AG and SR are supported by the Korean Institute for Basic Science (IBS-R022-A2-2023). GF was supported by an EMBO Long-Term Fellowship (ALTF 1132-2018). EAM acknowledges grants from Cancer Research UK (C13474/A27826) and the Wellcome Trust (219475/Z/19/Z). Work in SEP lab is supported by the European Research Council (ERC-2018-CoG-818625). Work in CPP lab is supported by the MRC (grant number MC_UU_00007/15). This work was supported by the Medical Research Council, as part of United Kingdom Research and Innovation (also known as UK Research and Innovation) (MRC file reference number MC_U105192715). For the purpose of open access, the MRC Laboratory of Molecular Biology has applied a CC BY public copyright licence to any Author Accepted Manuscript version arising.

## Author Contributions

GS and JR conceived the project. FNC analysed ChIP-seq, ATAC-seq and mRNA-seq datasets under the supervision of JA. GF and LL performed the transgenerational gene silencing microscopy assay under the supervision of EAM. SR and AG analysed GFP::SPT-2 distribution and germline defects. SP performed SNAP-H3.3 experiments under the supervision of SEP. AA performed ATAC-seq, and AA and GS performed GFP::SPT-2 ChIP-seq. JLP and NBR performed the initial RNA-seq analysis. LS-P and CPP identified the *C. elegans* SPT-2 orthologue. ACD performed EMSA assays. PA performed molecular modelling of CeSPT-2. RT performed the cloning of human expression vectors. GS performed the remaining experiments, and GS and JR wrote the manuscript.

## Data Availability

All plasmids and antibodies generated in this study can be requested to the MRC PPU DSTT: https://mrcppureagents.dundee.ac.uk/reagents-from-papers/Rouse-SPT2-paper-1 All NGS datasets have been deposited on GEO with accession number GSE224802. Other materials generated in this work will be made available upon request.

## Materials and Methods

### Computational protein sequence analysis

Multiple sequence alignments were generated with the program T-Coffee using default parameters^61^, slightly refined manually and visualized with the Belvu program^62^. Profiles of the SPT2 evolutionarily conserved regions as hidden Markov models (HMMs) were generated using HMMer. Profile-based sequence searches were performed against the Uniref50 protein sequence database^63^ using HMMsearch^64,65^.

### Structural modelling and conservation analysis

The crystal structure of human SPT2 (Protein Data Bank [PDB] code 5BS7)^25^ was used as a search template to generate a structural homology model for *C. elegans* SPT-2 using the homology-model server SWISS-MODEL (swissmodel.expasy.org)^66,67^. A multiple protein sequence alignment was generated by MUSCLE (ebi.ac.uk/Tools/msa/muscle)^68^. UCSF Chimera^69^ was used to align the atomic models of human and *C. elegans* Set using the “MatchMaker” function. Protein sequence conservation was mapped onto the structural model and figures generated in Chimera.

### Recombinant protein purification

All recombinant SPT2 proteins were expressed in Rosetta 2(DE3)pLysS BL21 *E. coli* bacteria (Novagen, 71401). Plasmids were transformed into Rosetta BL21 cells according to the manufacturer’s instructions and transformed bacteria were grown in LB medium supplemented with 35 μg/ml chloramphenicol and 50μg/ml of kanamycin.

#### Full-length CeSPT-2 (wild type and HBM) and HsSPT2 (His_6_-tagged)

Expression of 2xFlag-SUMO-CeSPT-2-His_6_ or of 2xFlag-SUMO-HsSPT2-His_6_ was induced by growing bacteria overnight at 20°C with 200μM IPTG. After cell lysis (Lysis buffer: 50mM Tris-HCl pH 8.1, 500mM NaCl, 10mM imidazole pH 8, 10% glycerol, 0.1mM TCEP supplemented with protease inhibitors), lysates were incubated with NiNTA beads for 1 hour rotating top-down at 4°C, and subsequently washed with Lysis buffer. Proteins were eluted by addition of 500mM imidazole (Elution buffer: 50mM Tris-HCl pH 8.1, 500mM NaCl, 500mM imidazole pH 8, 10% glycerol, 0.1mM TCEP supplemented with protease inhibitors) and Flag M2 beads were immediately added to the eluate, and incubated for 2 hours rotating top-down at 4°C. The M2 beads were washed twice with Lysis buffer supplemented with 10mM MgCl_2_ and 2mM ATP to remove folding chaperones. Flag-tagged proteins were eluted twice by adding Lysis buffer containing 0.5mg/ml Flag peptide (MRC PPU Reagents and Services). The eluate was incubated with 200nM His_6_-Ulp1 (prepared as previously described^70^) rotating top-down for 1 hour at 4°C. The cleaved 2xFlag-SUMO tag was removed by re-binding the eluate on Flag M2 beads rotating top-down for 1 hour at 4°C and collecting the unbound fraction. Proteins were concentrated in an Amicon Ultra Centrifugal tube with a molecular weight cut-off of 30kDa and frozen in liquid nitrogen.

#### CeSPT-2 HBD (aa. 552-661) and HsSPT2 HBD (aa. 571-685) wild type and HBMs

Expression of His_14_-SUMO-CeSPT-2 HBD or His_14_-SUMO-HsSPT2 HBD was induced by growing bacteria overnight at 20°C with 400μM IPTG. After cell lysis (Lysis buffer: 20mM Tris-HCl pH 7.5, 200mM NaCl, 40mM imidazole pH 8, 0.1mM TCEP supplemented with protease inhibitors), lysates were incubated with NiNTA beads for 2 hours rotating top-down at 4°C and washed with Lysis buffer. Proteins were eluted by addition of 500mM imidazole (Elution buffer: 20mM Tris-HCl pH 7.5, 500mM NaCl, 500mM imidazole pH 8, 0.1mM TCEP supplemented with protease inhibitors). The eluate was incubated with 60μM His_6_-Ulp1 to cleave the His_14_-SUMO tag and dialysed overnight against Dialysis buffer (20mM Tris-HCl pH 7.5, 500mM NaCl, 0.1mM TCEP). The cleaved His_14_-SUMO tag was removed by re-binding the eluate on NiNTA beads and collecting the unbound fraction containing the SPT2 HBD. Proteins were concentrated in a Amicon Ultra Centrifugal tube with molecular weight cut-off of 3kDa and frozen in liquid nitrogen.

### Histone H3-H4 pull-down

Protein LoBind tubes were used at all steps. For each reaction, 10μl of Pierce NHS-activated magnetic beads (ThermoFisher, 88826) were prepared according to the manufacturer’s instructions. Briefly, the beads were washed with 1ml of cold 1mM HCl and then were resuspended in 1mL of Coupling Buffer (500mM NaCl, 200mM NaHCO_3_, pH 8.3). 20pmol of recombinant histone H3.1-H4 tetramer (NEB, M2509S) were immediately added to the tube. The reaction was incubated overnight at 4°C with top to bottom rotation. The following day, the beads were washed twice with 0.1M glycine (pH 2.0) and once with MilliQ water. 1mL of Quenching Buffer (500mM ethanolamine, 500mM NaCl pH 8.5) was added to the beads, followed by 2 hours incubation at 4°C. The beads were then washed once with MilliQ and twice with Binding Buffer (20mM Tris-HCl pH 8.0, 500mM NaCl, 0.1% Triton-X100, 0.1mM TCEP). 5pmol of recombinant SPT2 proteins were added to the beads in 800μl Binding Buffer and incubated for 1.5 hours at 4°C, rotating top-down. The beads were then washed 3 times with Washing Buffer (20mM Tris-HCl pH 8.0, 700mM NaCl, 0.1% Triton-X100, 0.1mM TCEP), each time rotating for 15 minutes on a top-down wheel at 4°C. The beads were then boiled for 10 minutes in 1XLDS buffer (supplemented with 150mM DTT) and the supernatant was loaded on a 4-12% NuPage Bis-Tris gel. For the H3-H4 pull-down using the isolated HBD, the protocol was the same as above, except for the following differences: NHS-activated Sepharose 4 Fast Flow beads (Cytiva, 17090601) instead of magnetic beads were used; 1nmol of recombinant *Xenopus laevis* H3(Δ1-40 aa.)-H4 (a gift from Ramasubramanian Sundaramoorthy and Tom Owen-Hughes) and 0.5nmol of recombinant CeSPT-2 or HsSPT2 HBDs were used; the eluted material was run on a home-made 18% acrylamide gel and then stained with InstantBlue (Abcam) Coomassie protein staining.

### Electrophoretic Mobility Shift Assay

#### Preparation of labelled HJ substrates

An equimolar mixture of all four oligonucleotides (J3b(40), J3h(40), J3r(40) and J3x(40)) was 5’-^32^P-labelled, then annealed by slow-cooling. The four-way junction was purified by electrophoresis on a native 8% polyacrylamide gel and recovered by the crush and soak method followed by ethanol precipitation.

#### Binding assays

The substrate (10nM) was incubated with increasing amounts of SPT2 at 37°C for 15 minutes in binding buffer (25mM Tris-HCl pH 8.0, 55mM NaCl, 1mM EDTA, 1mM DTT, 0.1mg/ml BSA, 1% glycerol), then mixed with loading buffer (Ficoll 400, 2.5% final) and run on a native 8% polyacrylamide gel at 8 V/cm for 2 hours. The gel was then dried, exposed to a Storage Phosphor screen, and quantified with a Typhoon FLA 9500 phosphorimager (GE Healthcare).

Oligonucleotide sequences (5’ – 3’) are:

J3b(40): AGGGATCCGTCCTAGCAAGGGGCTGCTACCGGAAGCTTAC

J3h(40): GTAAGCTTCCGGTAGCAGCCTGAGCGGTGGTTGAATTCAC

J3r(40): GTGAATTCAACCACCGCTCAACTCAACTGCAGTCTAGAAC

J3x(40): GTTCTAGACTGCAGTTGAGTCCTTGCTAGGACGGATCCCT

### General *C. elegans* maintenance and strains

The list of strains used in this study is provided below. Unless otherwise indicated, worms were grown on 1xNGM media plates (3 g/L NaCl, 2.5 g/L peptone, 20 g/L agar, 5 mg/L cholesterol, 1mM CaCl_2_, 1mM MgSO_4_, 2.7g/L KH_2_PO_4_, 0.89 g/L K_2_HPO_4_) seeded with OP50 bacteria. Worms were routinely kept at 20°C. All *spt-2* alleles used in this study were generated by SunyBiotech using the CRISPR-Cas9 technology. All *spt-2* mutant strains were back-crossed 6 times against the reference wild type N2 strain.

### Analysis of GFP::SPT-2 tissue expression

For the analysis of the tissue expression of GFP::SPT-2, L4 animals were mounted in M9 buffer with 0.25mM levamisole on 2% agarose pads. Imaging was performed at 20°C using a microscope equipped with a 63X 1.25 NA oil lens (Imager M2; Carl Zeiss, Inc.) and a charge-coupled device camera (Axiocam 503 mono). Nomarski and GFP Z-stacks (2μm/slice) were sequentially acquired using the Zeiss acquisition software (ZEN 3.1 blue edition). The same LED intensity and acquisition time was used for all images (Nomarski (50%, 20ms), GFP (50%, 100ms)).

### Brood size analysis

Worms at the L4 stage were singled onto 1xNGM plates with a thin layer of OP50 bacteria and allowed to grow at either 20°C or 25°C. Worms were passaged to a new plate every 12-24 hours to keep generations separated, and fertilised eggs and L1 larvae were counted until each adult was not laying eggs anymore. Worms that were not alive by the end of the experiment were discarded from the analysis.

### Transgenerational sterility assay

Worms of the indicated genotypes were grown at 20°C before shifting to 25°C at the L4 stage. Either one or three L4 larvae (as indicated in the figure legend) were placed on each 1xNGM plate seeded with OP50 bacteria and grown at 25°C. Every 3 days, three L3-L4 larvae were transferred to a new plate. Worms were considered sterile when no progeny was found on the plate. Ten plates per genotype were used in each repetition of the assay.

### *hira-1* RNA interference

RNAi was performed by feeding worms with HT115 bacteria containing L4440 plasmids that express double-stranded RNA. The plasmid expressing *hira-1* dsRNA was obtained from a commercial RNAi library (clone III-2P01, Source Bioscience), and the empty L4440 vector was used as a control. dsRNA expression was induced by adding IPTG to a final 3mM concentration in LB media supplemented with 50μg/ml ampicillin, and RNAi bacteria were seeded on 1xNGM plates at OD_600_ equal 1. When the plate was dry, six L4 worms of the indicated genotypes were added to the plate and allowed to grow at 25°C until the next generation (2 days). New *hira-1* RNAi, or vector RNAi, plates were prepared as described above. Forty L2-L3 worms per genotype were singled on the RNAi plates and allowed to grow at 25°C for four/five days, after which plates were blindly scored for Pvl phenotype and presence of burst worms.

### Transgenerational memory inheritance

Three L4 larvae per genotype were plated on *gfp* RNAi-expressing bacteria (5 replicates for each *spt-2*^KO^ strain, or 3 replicates for each *spt-2*^HBM^ sibling) or empty vector L4440 bacteria (3 replicates). Generation 1 (G1) animals were analysed under a fluorescence microscope, and one silenced animal per replicate per genotype was plated onto HB101 bacteria. At each generation, one silenced animal was singled from each plate to produce the next generation, and the remaining adult progeny was analysed under a Kramer FBS10 fluorescence microscope. Animals were collected in M9 buffer, washed twice, quickly fixed in 70% ethanol, and deposited onto a glass slide coated with a 2% agarose pad. At least 35 animals per replicate per genotype were counted at each generation. Germline nuclear GFP brightness was scored as ‘on’ or ‘off’ by visual inspection (dim GFP expression was scored as ‘off’). Representative images were taken on a SP8 confocal fluorescence microscope (Leica) at 40X magnification.

### GFP::CeSPT-2 chromatin-immunoprecipitation and sequencing

Animals of the indicated genotype (grown at 20°C) were bleached, and embryos allowed to hatch overnight in M9 buffer. Approximately 300,000 worms were used per condition, divided in six 15cm plates seeded with thick HB101 bacteria. Synchronized adult animals (70h post-seeding) were washed 4 times in M9. Worms were flash-frozen in liquid nitrogen to obtain ‘popcorns’ which were ground using a BioPulverizer (MM400 Mixer Mill, Retsch) liquid nitrogen mill, until adult worms were broken in 3-4 pieces each. Worm powder was resuspended in cold PBS (supplemented with Roche cOmplete protease inhibitors). Freshly prepared EGS solution (ethylene glycol bis(succinimidyl succinate)) was added to a final 1.5mM concentration and incubated rotating for 8 minutes, followed by crosslinking with 1% methanol-free formaldehyde (ThermoFisher, 28908) for 8 minutes and quenching with glycine (final concentration 125mM). The crosslinked chromatin was then washed twice in PBS (with protease inhibitors) and once in FA buffer (50mM HEPES/KOH pH 7.5, 1mM EDTA, 1% Triton-X100, 0.1% sodium deoxycholate, 150mM NaCl) supplemented with protease (Roche Complete) and phosphatase (PhosStop) inhibitors. Chromatin was then sonicated using a Bioruptor (Diagenode) on High mode, 30 cycles at 30 sec ON, 30 sec OFF. A 30μl aliquot was de-crosslinked to confirm enrichment of DNA fragments between 100bp and 300bp. To de-crosslink, chromatin was spun down at maximum speed and the supernatant was transferred to a new tube and resuspended in FA buffer supplemented with 2μl of 10mg/ml RNase A (incubation at 37°C for 10 minutes) followed by Proteinase K treatment (incubation at 65°C for 1 hour). DNA concentration was measured using the Qubit assay (Qubit dsDNA High Sensitivity assay, ThermoFisher). The ChIP reaction was assembled as follows: 20μg of DNA were used per ChIP reaction, together with 10μg of GFP antibody (ab260) in 1mL of FA buffer (with protease and phosphatase inhibitors) supplemented with 1% sarkosyl. ChIP reactions were incubated overnight rotating at 4°C. For each reaction, 40μl of protein A magnetic beads slurry were blocked overnight in 1mL of FA buffer (with protease and phosphatase inhibitors) supplemented with 10μl of tRNA (Merck, R5636). The following day, beads were added to the IP reaction and incubated for a further 2 hours rotating at 4°C. The beads were then washed with 1ml of the following buffers (ice cold), each time rotating 5 minutes at 4°C: two times with FA buffer (with protease and phosphatase inhibitors); once with FA buffer supplemented to 500mM NaCl; once with FA buffer supplemented to 1M NaCl; once with TEL buffer (10mM Tris-HCl pH 8, 250mM LiCl, 1% NP-40, 1% sodium deoxycholate, 1mM EDTA) and finally twice in TE buffer (pH 8). DNA was eluted twice with 60μl of ChIP Elution buffer (1% SDS, 250mM NaCl in TE buffer pH 8), each time incubating for 15 minutes at 65°C (300rpm shaking). Eluted samples were treated with 2μl of 10mg/ml RNase A for 1 hour at 37°C and then de-crosslinked overnight at 65°C with 1.5μl of 20mg/mL Proteinase K. DNA was purified with PCR purification columns and DNA concentration was measured with the Qubit assay.

#### Sequencing library preparation

The library preparation was performed as previously described^71^ using a modified Illumina TruSeq protocol. Briefly, DNA fragments were first treated with End repair enzyme mix (NEB, E5060) for 30 min at 20°C in 50μl volume, and DNA fragments were subsequently recovered with one volume of AMPure XP beads (Beckman Coulter, A63880) mixed with one volume of 30% of PEG_8000_ in 1.25M NaCl. DNA was eluted in 17μl of water and further 3’ A-tailed using 2.5 units of Klenow 3’ to 5’ exo(-) (NEB, cat M0212) in 1X NEB buffer 2 supplemented with 0.2 mM dATP for 30 minutes at 37°C. Illumina Truseq adaptors were ligated to the DNA fragments by adding 25μl of 2X Quick Ligase buffer (NEB, M2200), 1μl of adaptors (1μl of a 1:250 dilution of Illumina stock solution), 2.5μl water and 1.5 ml of NEB Quick ligase (NEB, M2200). The reaction was incubated 20 minutes at room temperature and 5μl of 0.5M EDTA was added to inactivate the ligase enzyme. DNA was purified using 0.9 volume of AMPure XP beads, and DNA fragments were eluted in 20μl water. 1μl of DNA was used to set up a qPCR reaction to determine the number of cycles needed to get amplification to 50% of the plateau, which is the cycle number that will be used to amplify the library. 1μl of DNA was mixed with 5μl of KAPA Syber Fast 2X PCR master mix and 1μl of 25μM Truseq PCR Primer cocktail. Each library (19μl) was then amplified with 20μl of KAPA HiFi HotStart Ready Mix (KM2605) and subsequently size selected to a size between 250bp and 370bp. To achieve this, DNA was first purified with 0.7 volumes of AMPure beads to remove all fragments above 370bp, keeping the supernatant and discarding the beads. All DNA fragments were then recovered from the supernatant by adding 0.75 volumes of beads and 0.75 volumes of 30% PEG_8000_ in 1.25M NaCl, and eluted in 40μl of water. To recover fragments above 250bp, 0.8 volumes of beads were added to bind the library. DNA was then eluted in 10μl of water, quantified using a dsDNA HS Qubit kit, and the size distribution of the libraries was analyzed using an Agilent Tapestation.

### Processing of sequencing data

ATAC-seq and ChIP-seq reads were pre-processed using trim-galore (version 0.6.7; available at https://github.com/FelixKrueger/TrimGalore) and mapped on the *C. elegans* genome (wormbase release WS285) using bwa-mem (version 0.7.17)^72^. Reads with mapping quality (MAPQ) higher than 10 were extracted using Samtools^73^. ATAC-seq peaks were called by first generating read depth-normalized coverage tracks with MACS2^74^ (version 2.7.1; settings: --nomodel –extsize 150 –shift −75 --keep-dup all –SPMR -- nolambda) and then running YAPC^71^ (version 0.1) using bigwig tracks from both replicates for each condition as input. CeSPT-2 ChIP-seq peaks were called using MACS2 (settings: --SPMR --gsize ce --keep-dup all --nomodel --broad) using the GFP ChIP-seq samples from wild type animals as controls. The final SPT-2 peak set was defined by the strict intersection of the broad peaks called on each replicate.

RNA-seq data were aligned on annotated transcripts (Wormbase release WS285; gene types included: protein coding, lincRNAs and pseudogenes) using kallisto^75^ (version 0.45.0) to estimate gene expression (in TPM). Coverage profiles were obtained by aligning reads to the genome using STAR^76^ (version 2.7.1a).

### Annotation of CeSPT-2 bound genes

A gene was considered a CeSPT-2 target if more than 50% of the length of its longest annotated transcript was covered by a CeSPT-2 peak. The average CeSPT-2 coverage over CeSPT-2 targets was calculated using coverageBed from the BEDTools suite^77^ (v.2.30.0). Coverage plots over gene models were produced using the DeepTools suite^78^ (version 3.5.1).

### ATAC-seq

Animals of the indicated genotype (grown at 20°C) were bleached, and embryos allowed to hatch overnight in M9 buffer. Approximately 12,000 L1 worms were seeded onto a 15cm plates seeded with a thick lawn of HB101 bacteria. Gravid adult worms were collected 70h post-seeding and washed three times in M9 buffer. To assess that worm synchronization was equal between samples, 10μl of worm slurry was fixed in cold methanol overnight at − 20°C; the remaining slurry was frozen in ‘popcorns’. The methanol-fixed worms were stained with DAPI (final 1μg/ml) for 10 minutes, washes three times with PBS/0.1% Tween, and rehydrated overnight at 4°C in PBS/0.1% Tween. Worms were then mounted on a slide and inspected on a Leica SP8 UV microscope. ATAC-seq was performed as previously described^71^. Frozen worms (3-4 frozen popcorns) were broken by grinding in a mortar and pestle. The frozen powder was thawed in 10ml Egg buffer (25mM HEPES pH 7.3, 118mM NaCl, 48mM KCl, 2mM CaCl_2_, 2mM MgCl_2_) and washed twice with that buffer by spinning 2 minutes at 1500g. After the second wash, the pellet was resuspended into 1.5ml of Egg buffer containing 1mM DTT, protease inhibitors (Roche complete, EDTA free) and 0.025% of Igepal CA-630. Samples were dounced 20 times in a 7ml stainless tissue grinder (VWR) and then spun at 200g for 5min to pellet large debris. Supernatant containing nuclei was transferred into a new tube and nuclei were counted using a haemocytometer. One million nuclei were transferred to a new 1.5ml tube and spun at 1000g for 10 minutes, the supernatant was removed, and the nuclei resuspended in 47.5μl of Tagmentation buffer (containing 25μl of 2x Tagmentation buffer from Illumina and 22.5μl of water) before adding 2.5μl of Tn5 enzyme (Illumina Nextera kit) and incubated for 30 minutes at 37°C. Tagmented DNA was then purified using MinElute column (Qiagen) and amplified 10 cycles using the Nextera kit protocol. Amplified libraries were finally size selected using AMPure beads to recover fragments between 150 and 500bp and sequenced.

### Analysis of differentially accessible genes (ATAC-seq)

We used DiffBind^79^ (version 3.8.0) to identify genes showing differential accessibility in the *spt-2*^KO-A^ and the *spt-2*^HBM^ strains compared to the wild type. For this analysis, we had to redefine gene coordinates to avoid overlaps, which would be otherwise merged in DiffBind. The boundaries of the longest transcript for each gene were trimmed to remove overlaps with neighbouring transcripts; if more than 50% of the length of the transcript was trimmed, the whole gene locus was removed from the analysis. Genes were defined as differentially accessible if the |LFC| > 0 and FDR < 0.05, and their enrichment evaluated at non-CeSPT-2 targets and at CeSPT-2 targets divided in 4 equally-sized groups based on their CeSPT-2 levels.

### Collection of adult worms for total RNA extraction

Animals of the indicated genotype (grown at 20°C) were bleached, and embryos were allowed to hatch overnight in M9 buffer. Approximately 2000 L1 worms were seeded onto 10cm plates seeded with HB101 bacteria. Gravid adult worms were collected 70h post-seeding, washed three times in M9 buffer and the worm pellet was resuspended in 1mL Trizol (ThermoFisher, 15596026). This mixture was quickly thawed in a water bath at 37°C, followed by vortexing and freezing in liquid nitrogen; five freeze/thaw cycles were performed. Then, 200μl of chloroform were added, and the mixture was vortexed for 15 seconds and incubated at room temperature for 3 minutes. The mixture was spun down for 15 minutes at 15,000g in a refrigerated centrifuge (4°C). The upper aqueous phase was carefully moved to a new 1.5ml tube and RNA was retrieved by isopropanol precipitation. Briefly, 1μl of glycogen and 500μl of isopropanol was added and, after vortexing, the mixture was incubated overnight at −20°C. The next day, the mix was centrifuged for 15 minutes at 15,000g, 4°C and the supernatant removed. 1mL of 70% EtOH was added, and the tube was again centrifuged for 5 minutes at 15,000g, 4°C. After carefully removing the supernatant, the RNA pellet was allowed to dry at room temperature for a maximum of 5 minutes and then resuspended in 40μl DEPC-treated water (Invitrogen). The RNA quality was assessed on an Agilent Tapestation, using the RNA ScreenTape kit according to the manufacturer’s instructions (Agilent, 5067-5576 and 5067-5577).

### Poly(A) enrichment, library preparation and sequencing

Total RNA extracted from adult worms was treated with Turbo DNA-free kit (Invitrogen, AM1907) according to the manufacturer’s protocol. RNA concentration was measured with a Qubit instrument and 1μg of total RNA was used to make libraries. Poly(A) tail selection was performed using NEBNext Poly(A) mRNA Magnetic Isolation Module (NEB, E7490S) and libraries for next-generation sequencing were prepared using the NEBNext Ultra(tm) II Directional RNA Library Prep kit with Sample Purification Beads (NEB, E7765S). Libraries were indexed using NEBNext Multiplex Oligos for Illumina Index Primers Set 1 and 2 (E7335S and E7500S). Sequencing was performed on a NovaSeq instrument, 100bp paired end sequencing.

### Differential gene expression

We used DESeq2^80^ (version 1.34.0) to identify genes differentially expressed in the *spt-2*^KO^ and the *spt-2*^*HBM*^ strains compared to the wild type. Genes were defined as differentially expressed if the |LFC| > 0 and p.adj < 0.001. Volcano plots were generated using the EnhancedVolcano R package (available at https://github.com/kevinblighe/EnhancedVolcano). GO-enrichment analysis of differentially expressed genes was performed using clusterProfiler^81^. The overlap between genes upregulated in *spt-2* and *hira-1* mutant^36^ was tested using an hypergeometric test.

### RT-qPCR

For the retro-transcription reaction, 1μl of 50μM random hexamers and 1μl of 10μM dNTPs were added to 200-500ng of RNA in a 10μl volume reaction. The reactions were heated for 5 minutes at 65°C to denature secondary structures and allow random primer annealing. Then, 4μl of 5x First strand buffer, 2μl of 0.1M DTT, 1μl of RNaseOUT and 1μl of SuperScript RT III were added. This mix was incubated on a thermocycler with the following program: 10 minutes at 25°C, 1 hour at 50°C and 15 min at 70°C. Before proceeding with the qPCR reaction, the mix was diluted 1:5 with Sigma water.

qPCR primer sequences (5’ – 3’) are:

GSA144 gfp_ex1_F: GTGAAGGTGATGCAACATACGG (from ref. ^82^)

GSA145 gfp_ex1/2junction_R: ACAAGTGTTGGCCATGGAAC (from ref. ^82^)

GSA146 cdc-42 F: CTGCTGGACAGGAAGATTACG (from ref. ^82^)

GSA147 cdc-42 R: CTCGGACATTCTCGAATGAAG (from ref. ^82^)

GSA213 R06C1.4 F1: GTGTACGATCGTGAAACCGG

GSA214 R06C1.4 R1: CGAAGGTTGCGTCCATTGAA

GSA215 Y57G11B.5 F1: GCTTGTAATGCCGAGACGAG

GSA216 Y57G11B.5 R1: TGGAACATTTCGAGACGGGA

GSA217 asp-1 F1: GATTCCAGCCATTCGTCGAC

GSA218 asp-1 R1: TGATCCGGTGTCAAGAACGA

GSA221 clec-66 F1: TGCCATGACTAAATTCGCCG

GSA222 clec-66 R1: ACGCTCTCTTCTGTTGGTCA

GSA223 F08B12.4 F1: GAAAAGCGTCTTGGAAGGGG

GSA224 F08B12.4 R1: TTACTGGTGGTTTTGCTCGC

GSA225 C14C6.5 F1: CTACGACAATGGCACCAACC

GSA227 C14C6.5 R1: TTCATTCCTGGGCAGTCACT

GSA227 skr-10 F1: TGAGAGAGCTGCAAAGGAGA

GSA228 skr-10 R1: TGGAAGTCGATGGTTCAGCT

GSA350 H3.3-SNAP F2: CCTGGCTCAACGCCTACTTT (from ref. ^59^)

GSA351 H3.3-SNAP R2: GGTAGCTGATGACCTCTCCG (from ref. ^59^)

GSA352 H3.3-SNAP F3: TCGGAGAGGTCATCAGCTAC

GSA353 H3.3-SNAP R3: CAGAATGGGCACGGGATTTC

### Human cell culture

All cells were kept at 37°C under humidified conditions with 5% CO_2_. Parental U-2-OS cells were grown in DMEM (Life Technologies) supplemented with 100 U/ml penicillin, 100μg/ml streptomycin, 1% L-glutamate (Invitrogen), and 10% foetal bovine serum. U-2-OS SNAP-H3.3 cells were described before^49^ and were grown in the medium described above supplemented with 100μg/ml G418 (Formedium, G4181S). To generate stable U-2-OS Flp-In Trex cell lines, hygromycin was used to select for the integration of GFP-SPT2 constructs at the Flp-In recombination sites. U-2-OS Flp-In Trex cells expressing GFP-tagged SPT2 were grown in the medium described above supplemented with 100μg/ml hygromycin and 10μg/ml blasticidin. Expression of GFP–SPT2 was induced by addition of 1 μg/ml of tetracycline for 24 hours. All cell lines tested negative for mycoplasma contamination.

### siRNA transfection

Twenty-four hours prior to transfection, 80,000 U-2-OS cells were seeded per 6-well dish. siRNA transfection was performed with RNAiMax reagent (Invitrogen) according to the manufacturer’s protocol and all siRNAs were used to a final concentration of 50nM. Cells were harvested 48 hours post-transfection. siRNA sequences (5’ – 3’) are the following: Universal siRNA control (Sigma, SIC001)

siSPT2#1: GACCTATGACCGCAGAAGA

siSPT2#2: GTTACAATGGGATTCCTAT

siSPT2#3: GAGAATTCCTTGAACGAAA

siHIRA#1 GAAGGACUCUCGUCUCAUG (from ref. ^83^)

### Soluble/chromatin fractionation

To analyse soluble and chromatin-bound proteins by western blotting, cells were washed in cold PBS, scraped, and centrifuged at 1,500g for 10 minutes at 4°C. The pellet was incubated on ice for 10 minutes in CSK buffer (10mM PIPES pH 7, 100mM NaCl, 300mM sucrose, 3mM MgCl_2_)/0.5% Triton X-100, supplemented with Roche Complete Protease Inhibitor cocktail and PhosSTOP Roche phosphatase inhibitors, followed by 5 minutes centrifugation at 1,500g, 4°C (Soluble fraction). The pellet was washed with CSK/0.5% Triton, prior to resuspending in 1xLDS buffer supplemented with 10mM MgCl_2_, 150mM DTT and 250 units of Pierce Universal Nuclease (ThermoFisher, 88702), and incubated at 37°C shaking for 1 hour (Chromatin fraction).

### RNA extraction from human cells

RNA was extracted from U-2-OS cells according to the Qiagen RNeasy Mini Extraction kit. DNaseI digestion was performed on columns using the Qiagen RNase-Free DNase Set.

### Fluorescence analysis of human GFP-SPT2

To analyse GFP-SPT2 intensity on chromatin, U-2-OS Flp-In cells grown in 96-well plates (Greiner) were treated with 1μg/ml tetracycline for 24 hours to induce expression of GFP-tagged SPT2. Cells were then either directly fixed with 4% paraformaldehyde (10 minutes at room temperature), or pre-extracted for 5 minutes on ice with CSK/0.5% Triton buffer (supplemented with Roche Complete Protease inhibitor and PhosSTOP Phosphatase inhibitors) to remove soluble proteins, prior to fixation for 10 minutes with 4% paraformaldehyde at room temperature. DNA was stained with DAPI, and images were acquired using a Perkin Elmer Operetta high-content automated microscope (equipped with a 20X dry objective) and analysed using the Perkin Elmer Columbus software.

### Western blotting and antibodies

Primary antibodies: human SPT2 (in-house produced, DA010), GAPDH (Cell Signalling, clone 14C10, 2118S), GFP (Abcam, ab290), RPB1 (Cell Signalling, clone D8L4Y, 14958S), HIRA (Active Motif, 39457), His_6_ (Abcam, ab18184), H3 (Abcam, ab1791). Secondary IRDye LI-COR antibodies were used. Images were acquired using an Odyssey CLx LI-COR scanner.

### Human SPT2 antibody production

Polyclonal SPT2 antibodies were raised in sheep by the MRC PPU Reagents and Services Unit (University of Dundee) and purified against the SPT2 antigen aa. 385-685 (after depleting antibodies recognizing the epitope tags). Sheep DA010, 3^rd^ bleed, was used in this study. Sheep were immunised with the antigens followed by four further injections 28 days apart. Bleeds were performed seven days after each injection.

### SNAP-tag labelling

For labeling newly synthesized SNAP-tagged histones in U-2-OS cells stably expressing H3.3-SNAP, pre-existing SNAP-tagged histones were first quenched by incubating cells with 10 μM of the non-fluorescent SNAP reagent (SNAP-cell Block, New England Biolabs) for 30 min at 37°C followed by a 30 min wash in fresh medium and a 2-hour chase. The SNAP-tagged histones neosynthesized during the chase time were then pulse-labelled by incubating cells with 2 μM of the red-fluorescent SNAP reagent SNAP-cell TMR star (New England Biolabs) for 15 min at 37°C, followed by a 45 min wash in fresh medium. Cells were then directly fixed in 2% paraformaldehyde or pre-extracted before fixation in CSK buffer containing 0.5% Triton X-100. Samples were observed with a Leica DMI6000 epifluorescence microscope using a Plan-Apochromat 40x/1.3 oil objective. Images were captured using a CCD camera (Photometrics) and Metamorph software. Fiji software was used for image analyses using custom macros. Nuclei were delineated based on DAPI staining.

### Statistical analysis

The difference of variance between two populations was measured using Prism 9 software, and a Welch’s correction was applied when the variances were not equal. p values are provided and defined in the legend of the figure. The multiple comparisons in Fig. 5d and e were performed with One-way ANOVA test with Bonferroni’s correction. The overlap between genes upregulate in *spt-2* and *hira-1* mutant worms (Fig. 5g) was performed using a hypergeometric test.

### *C. elegans* strains

| Genotype | Strain name | Source |
| --- | --- | --- |
| WT N2 |  | CGC |
| *spt-2(syb1268) IV* | JRG30 | This work, from SunyBiotech |
| *spt-2(syb1269) IV* | JRG31 | This work, from SunyBiotech |
| *spt-2(syb2412[M627A]) IV* | JRG48 | This work, from SunyBiotech |
| *spt-2(syb1735[mAID-gfp::spt-2(wt)]) IV* | JRG44 | This work, from SunyBiotech |
| *spt-2(syb2435[mAID-gfp::spt-2(M627A)]) IV* | JRG45 | This work, from SunyBiotech |
| *spt-2(syb2412,syb4133[A627M]) IV* | PHX4133 | This work, from SunyBiotech |
| *mjls31 II* | SX461 | From ref. <sup>42</sup> |
| *mjls31 II; hrde-1(tm1200) III* | SX3448 | From ref. <sup>82</sup> |
| *mjls31 II; spt-2(syb1268) IV* | JRG49 | This work |
| *mjls31 II; spt-2(syb1269) IV* | JRG50 | This work |
| *mjls31 II; spt-2(syb2412[M627A]) IV* sibling 1 | JRG51 | This work |
| *mjls31 II; spt-2(syb2412[M627A]) IV* sibling 2 | JRG52 | This work |

### Plasmids

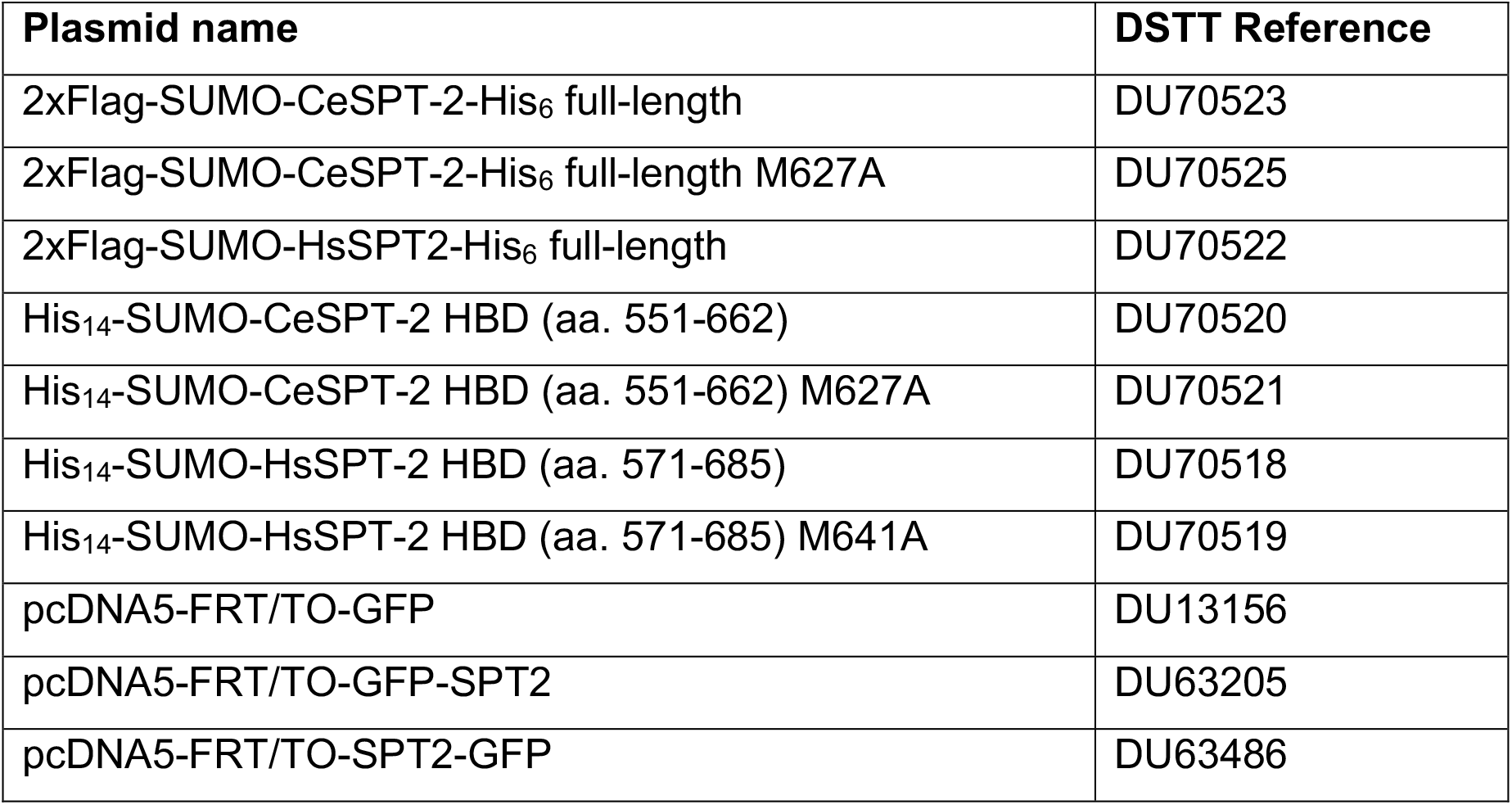
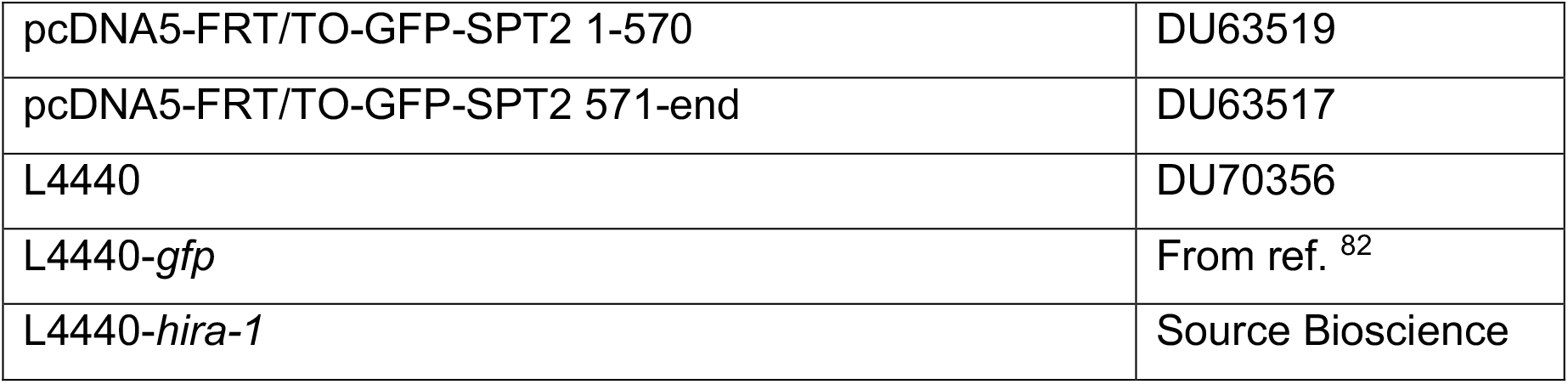

