## Supplementary Figures for "The histone chaperone activity of SPT2 controls chromatin structure and function in Metazoa"

**Figure S1**

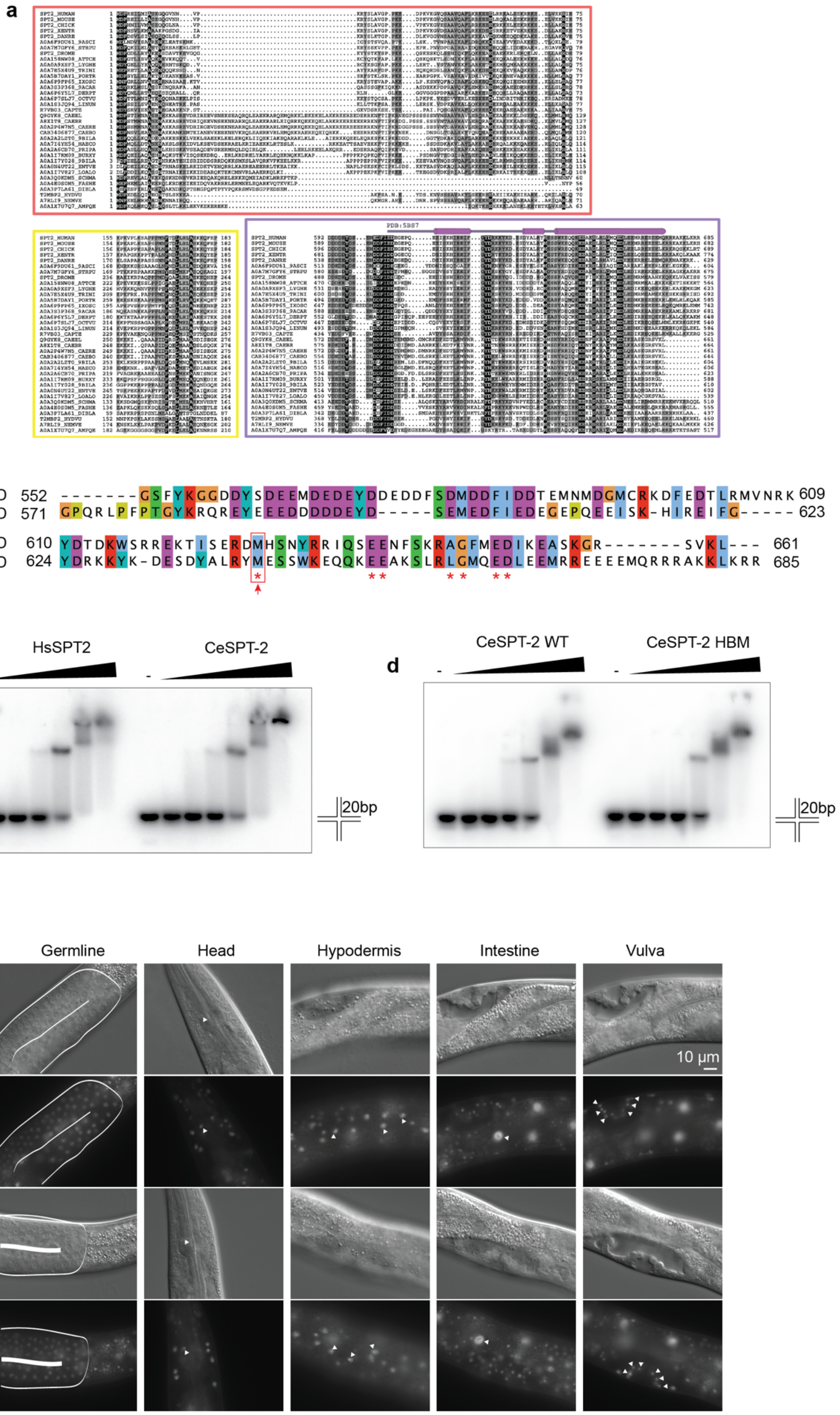
a
PDB:5BS7
b
CeSPT-2 HBD 552 -------GSFYKGGDDYSDEEMDEDEYDDEDDFSDMDDFIDDTEMNMDGMCRKDFEDTLRMVNRK 609
HsSPT-2 HBD 571 GPQRLPFPTGYKRQREYEEEDDDDDEYD-----SEMEDFIEDEGEPQEEISK-HIREIFG----- 623
CeSPT-2 HBD 610 YDTDKWSRREKTISERDMHSNYRRIQSEENFSKRAGFMEDIKEASKGR-------SVKL--- 661
HsSPT-2 HBD 624 YDRKKYK-DESDYALRYMESSWKEQQKEEAKSLRLGMQEDLEEMRREEEEMQRRRAKKLKRR 685
c
HsSPT2
CeSPT-2
20bp
d
CeSPT-2 WT
CeSPT-2 HBM
20bp
e
Germline
Head
Hypodermis
Intestine
Vulva
DIC
GFP::SPT-2 WT
10 µm
DIC
GFP::SPT-2 HBM

**Figure S1. Conservation and biochemical analysis of CeSPT-2.**

**a**, Multiple protein sequence alignments corresponding to the three conserved regions found in metazoan SPT2 proteins are shown inside coloured boxes in red, yellow, and violet, respectively. Above the alignment that corresponds to the conserved SPT2 C-terminal region (in violet), the secondary structure of the experimentally determined H3-H4 HDB of human SPT2 (PDB: 5BS7[25]) is shown (cylinders correspond to alpha helices). The amino acid colouring scheme indicates the average BLOSUM62 score (correlated to amino acid conservation) in each alignment column: black (greater than 3.5), grey (between 3.5 and 1.5) and light grey (between 1.5 and 0.5). Sequences are named according to their UniProt identifier. Species abbreviations: SPT2_HUMAN, *Homo sapiens*; SPT2_MOUSE, *Mus musculus*; SPT2_CHICK, *Gallus gallus*; SPT2_XENTR, *Xenopus tropicalis*; SPT2_DANRE, D*anio rerio*; A0A6F9DU61_9ASCI, *Phallusia mammillata*; A0A7M7GFY6_STRPU, *Strongylocentrotus purpuratus*; SPT2_DROME, *Drosophila melanogaster*; A0A158NW08_ATTCE, *Atta cephalotes*; A0A0A9XSF3_LYGHE, *Lygus hesperus*; A0A7E5X4U9_TRINI, *Trichoplusia ni*; A0A5B7DAY1_PORTR, *Portunus trituberculatus*; A0A6P9FP65_IXOSC, *Ixodes scapularis*; A0A3S3P368_9ACAR, *Dinothrombium tinctorium*; A0A6P6Y5L7_DERPT, *Dermatophagoides pteronyssinus*; A0A6P7SLJ7_OCTVU, *Octopus vulgaris*; A0A1S3JQ94_LINUN, *Lingula unguis*; R7VB03_CAPTE, *Capitella teleta*; Q9GYK8_CAEEL, *Caenorhabditis elegans*; A8X1T8_CAEBR, *Caenorhabditis briggsae*; A0A2P4W7N5_CAERE, *Caenorhabditis remanei*; CAB3406877_CAEBO, *Caenorhabditis bovis*; A0A2A2LZT0_9BILA, *Diploscapter pachys*; A0A7I4YH54_HAECO, *Haemonchus contortus*; A0A2A6CB70_PRIPA, *Pristionchus pacificus*; A0A1I7RM09_BURXY, *Bursaphelenchus xylophilus*; A0A1I7Y028_9BILA, *Steinernema glaseri*; A0A0N4UT22_ENTVE, *Enterobius vermicularis*; A0A1I7V827_LOALO, *Loa loa*; A0A3Q0KDM5_SCHMA, *Schistosoma mansoni*; A0A4E0S0M5_FASHE, *Fasciola hepatica*; A0A3P7LA61_DIBLA, *Dibothriocephalus latus*; T2MBP2_HYDVU, *Hydra vulgaris*; A7RLI9_NEMVE, *Nematostella vectensis*; A0A1X7U7Q7_AMPQE, *Amphimedon queenslandica*. **b,** Alignment of human and worm SPT2 histone binding domains. Red asterisks, residues in HsSPT2 reported to be required for the interaction with histones. Red arrow/box indicated the position of the Met residue histone binding defective mutation: M641 (human) / M627 (worm). **c, d**, Electrophoretic mobility shift assay (EMSA) showing binding of full-length recombinant CeSPT-2 or HsSPT2 (c), or of CeSPT-2 wild type and HBM (d), to synthetic cruciform DNA. Each arm of the four-way DNA junction contains 20 base pairs. For each panel, one representative experiment out of two is shown. **e,** Expression of GFP::CeSPT2 WT or HBM in the indicated tissues of L4 larvae. Images were taken with the same intensity and acquisition time.

### Figure S2

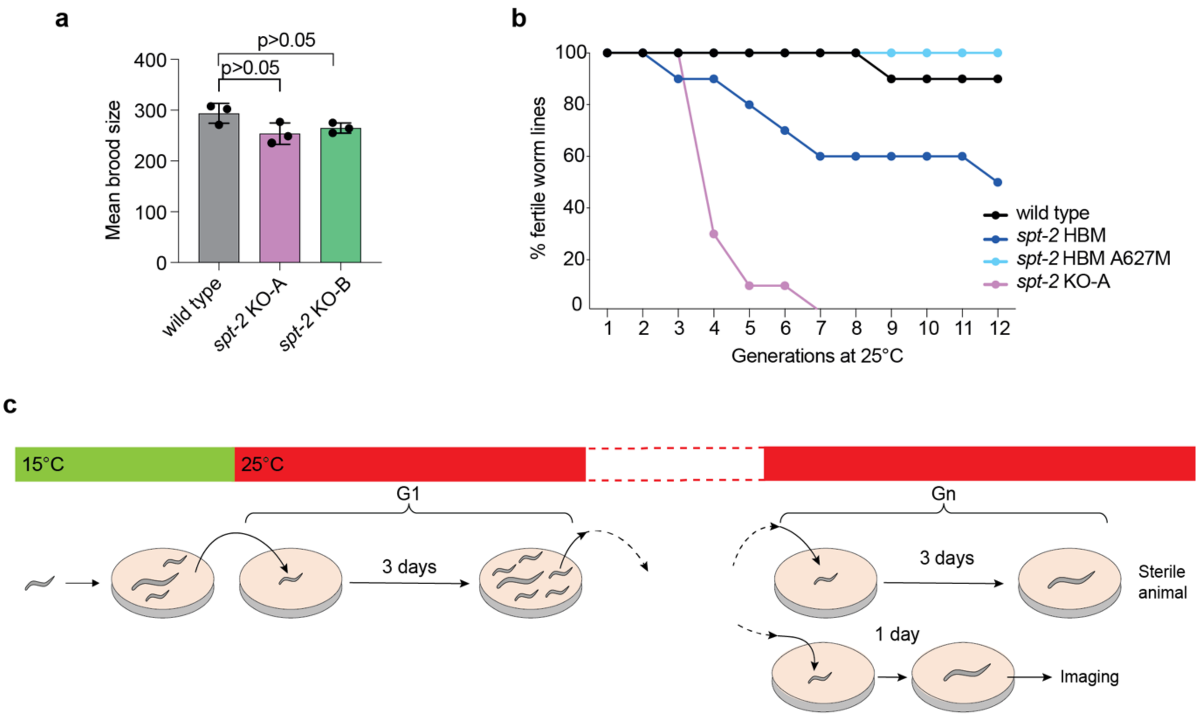


**d**

wild type

| Line | G1 | G2 | G3 | G4 | G5 |
|---|---|---|---|---|---|
| 1 | W | W | W | W* | W |
| 2 | W | W | W | W | W |
| 3 | W | W | W | W | W |
| 4 | W | W | W | W | W |
| 5 | W | W | W | W | W |
| 6 | W | W | W | W | S |
| 7 | W | W | W | W | M |

*spt-2* KO-B

| Line | G1 | G2 | G3 | G4 | G5 |
|---|---|---|---|---|---|
| 1 | W | W | W | W | W |
| 2 | W | W | W | W | W |
| 3 | W | W | W | W | W |
| 4 | W | W | W | W | S |
| 5 | W | W | W | F** | S*** |
| 6 | W | W | W | S | |
| 7 | W | W | W | W | M |

W: Wild type
M: Medium progeny
F: Few progeny
S: Sterile

* representative images shown in Fig. 2f
** representative images shown in Fig. 2g-j
*** representative images shown in Fig. 2k, S2e

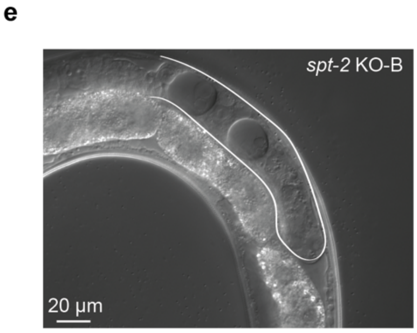

**Figure S2. Further characterization of *spt-2* mutant worms.**

**a**, Mean brood size of the indicated worm strains grown at 20°C. Worms at the L4 stage were singled onto plates and their total brood was counted. Each point represents the mean brood size of at least 5 worms, and the brood counting experiment was independently repeated three times. n=3; data are represented as mean ± S.D. **b**, Transgenerational sterility assay. Three L4 stage worms of the indicated genotypes were shifted to 25°C and grown at that temperature for the indicated number of generations. Every generation (3 days), three L3-L4 worms were moved to a new plate. A worm line was considered sterile when no progeny was found on the plate. Ten plates per genotype were used. **c, d**, Single wild type and *spt-2*$^{KO\text{-}B}$ L4 worms were shifted to 25°C and grown at that temperature for the indicated number of generations; every generation (3 days), one L4 worm was moved to a new plate. Siblings of the worms that, 3 days after the L4 stage, showed reduced or no progeny were subjected to microscopic observation 1 day after the L4 stage. The progeny size of each worm line was assessed as indicated (d). W, wild type: 200-250 worms. M, medium: 100 worms. F, few: less than 20 worms. S, sterile: no progeny. n=7 independent worm lines were used. **e**, Germline of a *spt-2*$^{KO\text{-}B}$ worm grown at 25°C showing vacuolization.

### Figure S3

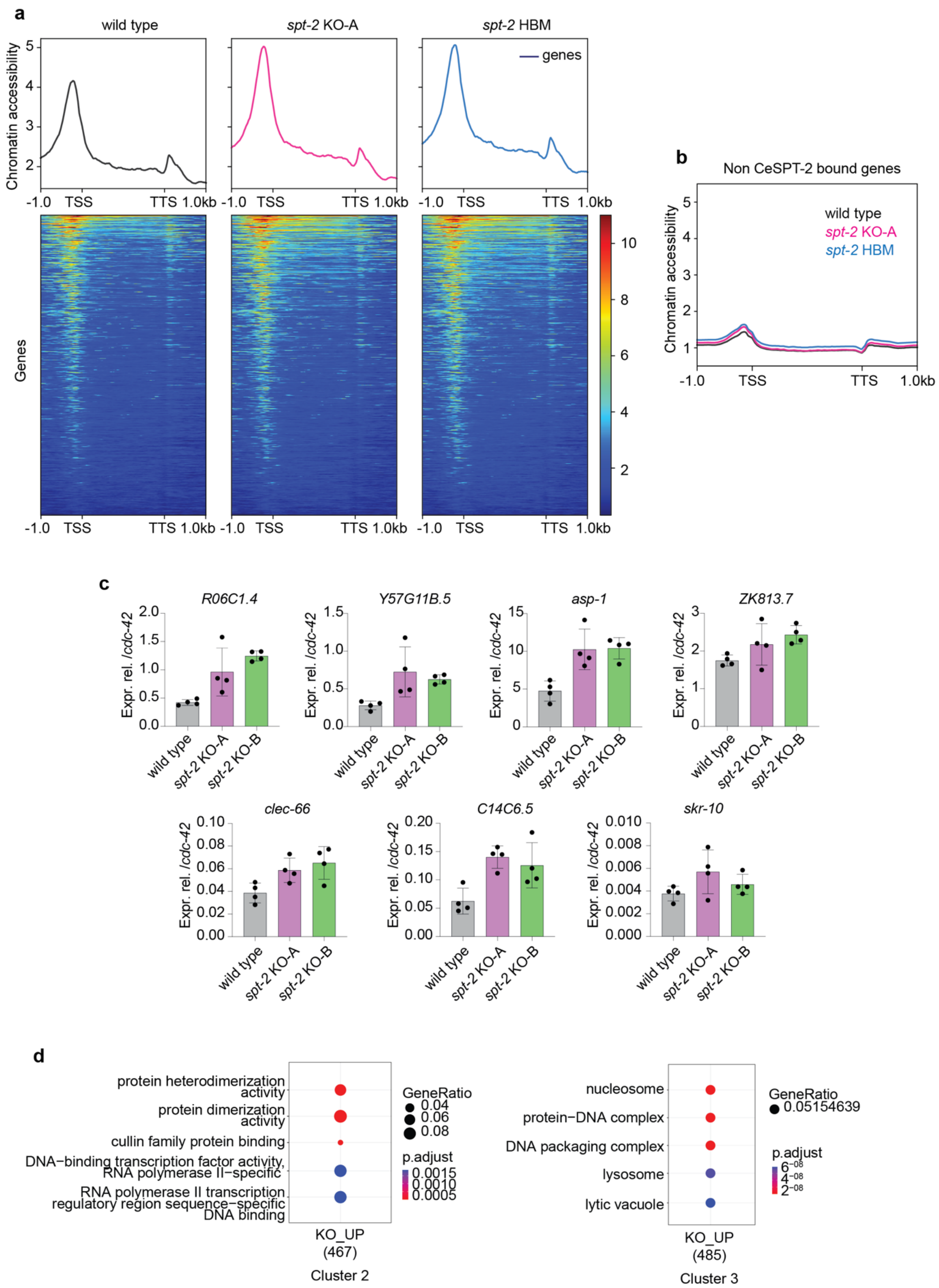

a
wild type
spt-2 KO-A
spt-2 HBM
genes
Chromatin accessibility
5
4
3
2
-1.0
TSS
TTS
1.0kb
Genes
10
8
6
4
2
b
Non CeSPT-2 bound genes
wild type
spt-2 KO-A
spt-2 HBM
Chromatin accessibility
c
R06C1.4
Y57G11B.5
asp-1
ZK813.7
clec-66
C14C6.5
skr-10
Expr. rel. /cdc-42
wild type
spt-2 KO-A
spt-2 KO-B
d
protein heterodimerization activity
protein dimerization activity
cullin family protein binding
DNA−binding transcription factor activity, RNA polymerase II−specific
RNA polymerase II transcription regulatory region sequence−specific DNA binding
GeneRatio
0.04
0.06
0.08
p.adjust
0.0015
0.0010
0.0005
KO_UP
(467)
Cluster 2
nucleosome
protein−DNA complex
DNA packaging complex
lysosome
lytic vacuole
GeneRatio
0.05154639
p.adjust
6−08
4−08
2−08
KO_UP
(485)
Cluster 3

**Figure S3. ATAC-seq and mRNA-seq analysis in spt-2 mutant worms.**
**a**, Heatmap of chromatin accessibility levels in the indicated strains, relative to Fig. 4d. **b**, Chromatin accessibility at CeSPT-2 non-target genes in *spt-2*$^{KO\text{-}A}$ and *spt-2*$^{HBM}$ adult worms, measured by ATAC-seq and expressed in reads per million. **c**, qPCR analysis for the indicated genes, relative to *cdc-42* expression. Each point indicates an independent replicate with worms harvested on different days, n=4. The RNA used for the qPCR was harvested independently from the RNA used for the mRNA-seq analysis. **d**, Gene Ontology analysis of the genes upregulated in *spt-2* null worms.

### Figure S4

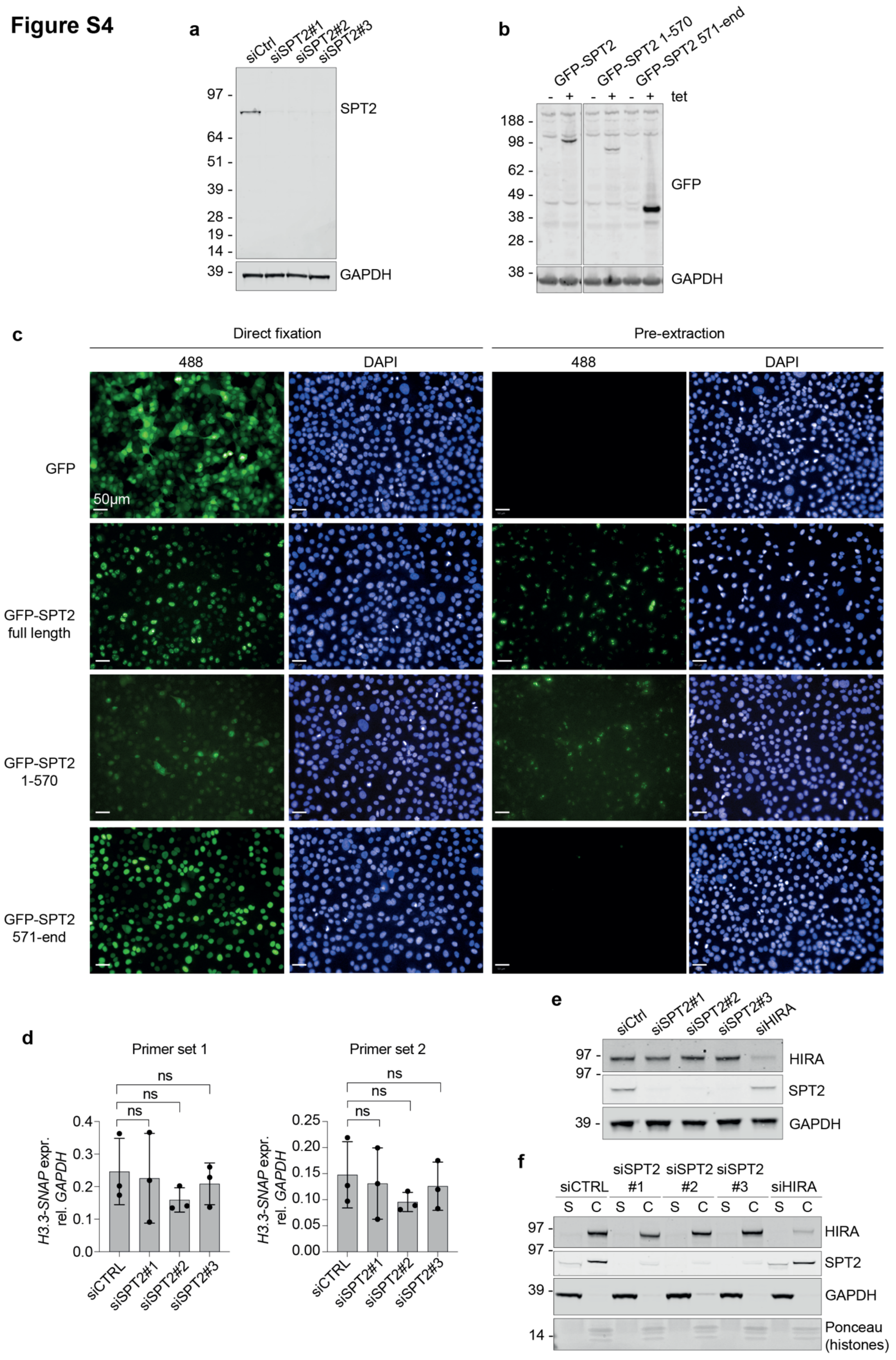
a
siCtrl
siSPT2#1
siSPT2#2
siSPT2#3
97 -
64 -
51 -
39 -
28 -
19 -
14 -
SPT2
39 -
GAPDH
b
GFP-SPT2
GFP-SPT2 1-570
GFP-SPT2 571-end
- + - + - + tet
188 -
98 -
62 -
49 -
38 -
28 -
GFP
38 -
GAPDH
c
Direct fixation
Pre-extraction
488
DAPI
488
DAPI
GFP
50μm
GFP-SPT2
full length
GFP-SPT2
1-570
GFP-SPT2
571-end
d
Primer set 1
ns
ns
ns
0.4
0.3
0.2
0.1
0.0
H3.3-SNAP expr.
rel. GAPDH
siCTRL
siSPT2#1
siSPT2#2
siSPT2#3
Primer set 2
ns
ns
ns
0.25
0.20
0.15
0.10
0.05
0.00
H3.3-SNAP expr.
rel. GAPDH
siCTRL
siSPT2#1
siSPT2#2
siSPT2#3
e
siCtrl
siSPT2#1
siSPT2#2
siSPT2#3
siHIRA
97 -
HIRA
97 -
SPT2
39 -
GAPDH
f
siCTRL
siSPT2
#1
siSPT2
#2
siSPT2
#3
siHIRA
S C S C S C S C S C
97 -
HIRA
97 -
SPT2
39 -
GAPDH
14 -
Ponceau
(histones)

76

**Figure S4. Validation of SPT2 antibodies and siRNAs.**

**a**, Western blot against HsSPT2 in total extracts of SPT2-depleted U-2-OS cells. **b**, Tetracycline-dependent expression of GFP-tagged HsSPT2 full length or the indicated truncation mutants in U-2-OS Flp-In cells. **c**, Representative high-content microscopy images relative to Fig. 5b are shown. Image contrast was applied equally to the directly fixed and pre-extracted samples of each cell line; contrast was instead adjusted independently for the different cell lines, which express different levels of GFP-tagged HsSPT2 (as shown in Fig. S4b). **d**, qPCR for the expression of H3.3-SNAP with two primer sets. Cells were harvested 48 hours post-transfection. n=3; data are represented as mean ± S.D. Primer set 1: $p= 0.8445$ (siCtrl vs siSPT2#1); 0.2364 (siCtrl vs siSPT2#2); 0.6159 (siCtrl vs siSPT2#3). Primer set 2: $p= 0.7704$ (siCtrl vs siSPT2#1); 0.2430 (siCtrl vs siSPT2#2); 0.6545 (siCtrl vs siSPT2#3). **e**, Total cell extracts of siRNA-treated U-2-OS cells. Cells were harvested 48 hours post-transfection. One representative experiment out of two is shown. **f**, U-2-OS cells were first depleted of HsSPT2, or HIRA, by siRNA treatment and soluble/chromatin fractions were obtained 48 hours post-transfection (S, soluble fraction and C, chromatin fraction). One representative experiment out of two is shown.

### Supplementary Table 1

**Replicate 1**

| | wild type | | | | | | | | | | | | | | |
|---|---|---|---|---|---|---|---|---|---|---|---|---|---|---|---|
| **Worm line** | 1 | 2 | 3 | 4 | 5 | 6 | 7 | 8 | 9 | 10 | 11 | 12 | 13 | 14 | 15 |
| **Brood size** | 138 | 123 | 153 | 155 | 171 | 161 | 153 | 147 | 113 | 109 | 125 | 104 | 150 | 98 | 137 |

| | *spt-2 KO-A* | | | | | | | | | | | | | | |
|---|---|---|---|---|---|---|---|---|---|---|---|---|---|---|---|
| **Worm line** | 1 | 2 | 3 | 4 | 5 | 6 | 7 | 8 | 9 | 10 | 11 | 12 | 13 | 14 | 15 |
| **Brood size** | 63 | 111 | dead | 97 | 0 | 0 | 171 | 109 | 122 | 65 | 80 | 0 | 0 | 24 | 0 |

| | *spt-2 KO-B* | | | | | | | | | | | | | | |
|---|---|---|---|---|---|---|---|---|---|---|---|---|---|---|---|
| **Worm line** | 1 | 2 | 3 | 4 | 5 | 6 | 7 | 8 | 9 | 10 | 11 | 12 | 13 | 14 | 15 |
| **Brood size** | 40 | 92 | 172 | 99 | 116 | 62 | 159 | 120 | 120 | 141 | 126 | 143 | 183 | 59 | 107 |

**Replicate 2**

| | wild type | | | | | | | | | |
|---|---|---|---|---|---|---|---|---|---|---|
| **Worm line** | 1 | 2 | 3 | 4 | 5 | 6 | 7 | 8 | 9 | 10 |
| **Brood size** | 260 | 176 | 194 | 232 | 155 | 205 | 150 | 185 | 199 | 211 |

| | *spt-2 KO-A* | | | | | | | | | | | | | | | | | | | |
|---|---|---|---|---|---|---|---|---|---|---|---|---|---|---|---|---|---|---|---|---|
| **Worm line** | 1 | 2 | 3 | 4 | 5 | 6 | 7 | 8 | 9 | 10 | 11 | 12 | 13 | 14 | 15 | 16 | 17 | 18 | 19 | 20 |
| **Brood size** | 212 | 196 | 189 | 120 | 109 | 201 | 192 | 192 | 175 | 214 | 148 | 49 | 221 | 220 | 223 | 180 | 233 | 129 | 195 | 192 |

| | *spt-2 KO-B* | | | | | | | | | | | | | | | | | | | |
|---|---|---|---|---|---|---|---|---|---|---|---|---|---|---|---|---|---|---|---|---|
| **Worm line** | 1 | 2 | 3 | 4 | 5 | 6 | 7 | 8 | 9 | 10 | 11 | 12 | 13 | 14 | 15 | 16 | 17 | 18 | 19 | 20 |
| **Brood size** | dead | 218 | 198 | 254 | 162 | 227 | 229 | 185 | 203 | 142 | 224 | 155 | 199 | 187 | 164 | 234 | 293 | 167 | 262 | 204 |

**Replicate 3**

| | wild type | | | | | | | | | |
|---|---|---|---|---|---|---|---|---|---|---|
| **Worm line** | 1 | 2 | 3 | 4 | 5 | 6 | 7 | 8 | 9 | 10 |
| **Brood size** | 181 | 155 | 247 | 192 | dead | 212 | 128 | 231 | 259 | 129 |

| | *spt-2 KO-A* | | | | | | | | | |
|---|---|---|---|---|---|---|---|---|---|---|
| **Worm line** | 1 | 2 | 3 | 4 | 5 | 6 | 7 | 8 | 9 | 10 |
| **Brood size** | 181 | dead | 16 | 190 | 151 | 185 | 189 | 147 | 180 | 174 |

| | *spt-2 KO-B* | | | | | | | | | |
|---|---|---|---|---|---|---|---|---|---|---|
| **Worm line** | 1 | 2 | 3 | 4 | 5 | 6 | 7 | 8 | 9 | 10 |
| **Brood size** | 77 | dead | dead | 112 | 98 | 128 | 199 | 121 | 105 | 22 |

**Supplementary Table 1. Brood size of *spt-2* null mutant worms grown at 25°C.** Worms at the L4 stage were singled onto plates at 25°C and their total brood was counted. The brood size of each worm line from three independent replicates is indicated. A worm was scored as 'dead' if it was not alive by the end of the assay.
